# Decoding Secondary Motor Cortex Neuronal Activity during Cocaine Self-Administration: Insights from Longitudinal *in vivo* Calcium Imaging

**DOI:** 10.1101/2024.05.20.594996

**Authors:** Yingying Chen, Haoying Fu, Amith Korada, Michal A. Lange, Chandrashekar Rayanki, Tao Lu, Dongbing Lai, Shiaofen Fang, Changyong Guo, Yao-Ying Ma

## Abstract

**Background:** We recently reported that the risk of cocaine relapse is linked to hyperexcitability in the secondary motor cortex (M2) after prolonged withdrawal following intravenous self-administration (IVSA). However, the neuronal mechanisms underlying drug-taking behaviors and the response of M2 neurons to contingent drug delivery remain poorly understood.

**Methods:** Mice received cocaine as reinforcement (RNF) following active lever presses (ALP) but not inactive lever presses (ILP). Using *in vivo* calcium imaging during cocaine IVSA, we tracked M2 neuronal activity with single-cell resolution. We then analyzed Ca^2+^ transients in M2 at the early *vs*. late stages during the 1-hr daily sessions on IVSA Day 1 and Day 5.

**Results:** M2 neurons adapted to both operant behaviors and drug exposure history. Specifically, saline mice showed a reduction in both saline taking behaviors and Ca^2+^ transient frequency with the 1-hr session. In contrast, cocaine mice maintained high ALP and RNF counts, with increased Ca^2+^ transient frequency and amplitude on Day 1, persisting through Day 5. Compared to saline controls, cocaine mice exhibited a lower % of positively responsive neurons (Pos) and higher % of negatively responsive neurons (Neg) before ALP and after RNF, a difference not seen before ILP. Furthermore, as drug-taking behaviors progressed during the daily session, cocaine mice showed greater neuronal engagement with a larger population, particularly linked to ALP and RNF, with reduced overlap in neurons associated with ILP.

**Conclusion:** The M2 undergoes dynamic neuronal adaptations during early drug-taking behaviors, indicating its role as a potential substrate mediating the persistence of drug-seeking behaviors in cocaine relapse.

## INTRODUCTION

Addiction is often attributed to conscious drug-seeking and taking behaviors. However, clinical observations of over 400 cocaine-dependent individuals reveal a different perspective. The most common self-reported reason for taking cocaine was impulsive action without a known cause (41%), rather than conscious craving (2%) (Miller and Gold, 1994). This suggests that drug-seeking and taking behaviors can become highly ritualized and automatic, frequently occurring without the individual’s awareness of any conscious thought or specific craving. We proposed that addiction is rooted in deeply ingrained motor patterns, likely mediated by the motor cortex. Early drug use and the establishment of addiction-related behaviors may represent the formation of drug-associated motor skills. Unlike typical motor skills, addiction-related motor patterns may develop with heightened efficiency due to the reinforcement (RNF) earned by drug-taking behavior (Willuhn and Steiner, 2009). Our recent study, the first to identify the secondary motor cortex (M2) as a novel region critically involved in addiction, demonstrated that hyperexcitability in M2 neurons heightens the risk of cocaine relapse after prolonged withdrawal (Huang and Ma, 2023a). This alteration in neuronal activity appears to be a consequence of initial cocaine experience. Therefore, we hypothesize that the initial experience of cocaine intake directly influences neuronal activity in M2, and understanding this effect is essential for unraveling the role of motor cortex in addiction.

The most reliable procedure for mimicking human SUD in laboratory animals is intravenous self-administration (IVSA) of the drug (Enga et al., 2016; Johanson, 1990; Johanson and Balster, 1978; Mayer-Blackwell et al., 2014; Zhang et al., 2014; Zhang et al., 2016). Traditional addiction research typically reports only the total number of IVSA events. These include active lever presses (ALP) that immediately trigger the delivery of reinforcers (e.g., cocaine IV infusion, denoted as RNF). It also accounts for inactive lever presses (ILP) which have no contingent consequence. Repeated drug taking behaviors during the IVSA session are experience-dependent. Each IVSA event (i.e., ALP, ILP, and RNF) has the potential to remodel neuronal function which, in turn, induces adaptations affecting the response to subsequent IVSA event. The interval between two consecutive IVSA events or between two IVSA sessions may serve as a critical consolidation period, allowing the brain to reorganize the experience-dependent changes and prepare for the upcoming behavioral demands. Thus, from the first, to the 10^th^ and even the 100^th^ IVSA event, the driving source from the brain and the effects on the brain may differ uniquely each time. Although addiction study has progressed unabatedly for decades, no studies have yet explored the dynamic changes in any brain region, including M2, at the neuronal level, before and after each of the drug taking behaviors.

Benefited from the advances in imaging technology and progress in fluorescent Ca^2+^ indicators(Cai et al., 2016; Ghosh et al., 2011; Zhang et al., 2019), *in vivo* Ca^2+^ imaging, which provides both high spatial and temporal resolution with high throughput in detecting individual neuronal activity, has recently been used in addiction research. However due to technical challenges related to Ca^2+^ imaging during IVSA (e.g., potential interference between the IV line for the IVSA procedure and the miniScope coax cable, head immobilization when multiphoton microscopy is used), the application of *in vivo* Ca^2+^ imaging recordings in addiction models has been limited to oral self-administration, passive injections, or head-fixed animals (Marchette et al., 2021; Sun and Giocomo, 2022; Vollmer et al., 2022). Considering the precise dosing and accurate timing of the IVSA and its high validity when animals are fully awake and freely moving (Colucci et al., 2014; Ma et al., 2014; Ma et al., 2016; Swain et al., 2021), we established a dual-channel commutator system (Doric) which allowed us to collect *in vivo* Ca^2+^ imaging data when the animals were allowed to operantly take drugs during the IVSA procedure. Together with our well-established pipeline (**see more details in MATERIALS AND METHODS**) in processing the Ca^2+^ imaging data, the current study was able to monitor cortical ensemble activity at the single cell level, in real time with high temporal solution and high throughput in each IVSA session, as well as longitudinally across multiple IVSA sessions. Specifically, five 1-hr daily IVSA sessions were used to model the initial cocaine taking behaviors. Besides high temporal resolution analyses of IVSA events (**Fig. 1**), the data from Ca^2+^ imaging recordings in M2 were analyzed by (1) comparing different time blocks (i.e., the first *vs*. the last 15 min) within a daily IVSA session and between different daily sessions (i.e., the first *vs*. the last IVSA day) (**Fig. 2**), (2) associating Ca^2+^ transients with an IVSA event (ALP, ILP, or RNF) (**Figs. 3-5, S1-6**), and (3) correlating Ca^2+^ transients with the cumulative effects of IVSA events (**Figs. 6-8, S7**).

**Figure 1.**
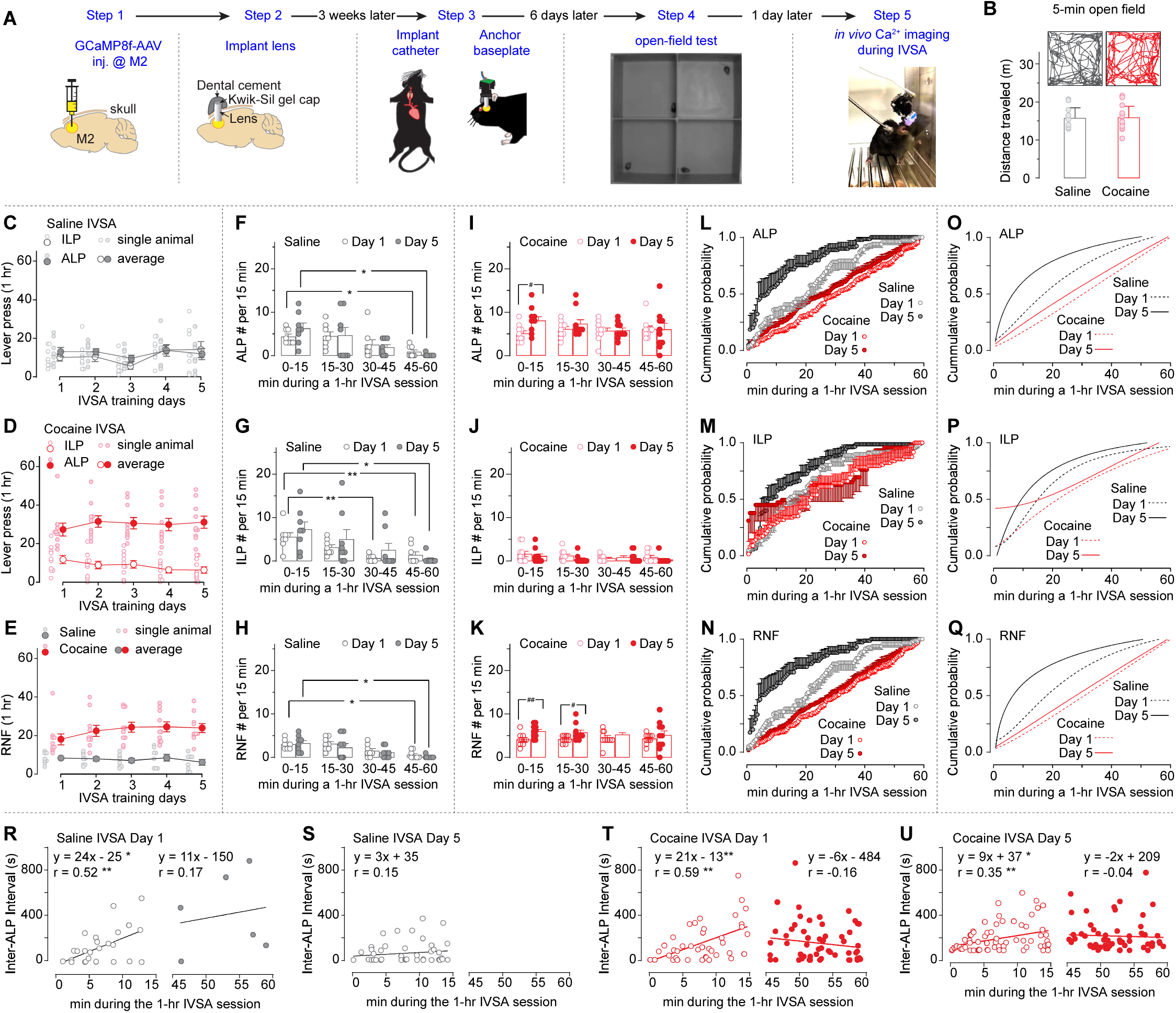
Cocaine taking behaviors. **A**, Experimental timeline **B**, No differences were observed in the total distance traveled during the 5-min open field test before the IVSA procedure between mice assigned to saline *vs*. cocaine treatment (t_16_=0.1, p=0.95). **C**, Individual mouse values (small circles) and summarized data (large circles) showing no differences between ALP *vs*. ILP during the 5 daily IVSA sessions in saline mice (ALP/ILP × IVSA day interaction F_4,28_=2.8, p=0.04; ALP/ILP F_1,7_=0.4, p=0.53; and IVSA day F_4,28_=0.8, p=0.53). **D**, Individual mouse values (small circles) and summarized data (large circles) showing significantly higher ALP, relative to ILP, during the 5 daily IVSA sessions in cocaine mice (ALP/ILP F_1,9_=55.6, p<0.01, in spite of ALP/ILP × IVSA day interaction F_4,36_=0.7, p=0.60; and IVSA day F_4,36_=2.0, p=0.11). **E**, Individual mouse values (small circles) and summarized data (large circles) showing a significantly higher RNF number in cocaine mice, relative to saline mice, during the 5 daily IVSA sessions (saline/cocaine × IVSA day interaction F_4,64_=3.1, p=0.02; saline/cocaine F_1,16_=29.7, p<0.01; and IVSA day F_4,64_=1.9, p=0.13). **F-H**, Individual mouse values (small circles) and summarized data (large circles) from saline mice showing gradual decreases in the numbers of ALP (**F**, 15-min slot F_2,14_=8.1, p<0.01, despite of IVSA day × 15-min slot interaction F_2,14_=1.6, p=0.24, IVSA day F_1,7_<0.1, p=0.82), ILP (**G**, 15-min slot F_2,14_=10.9, p<0.01, despite of IVSA day × 15-min slot interaction F_2,13_=0.9, p=0.41, IVSA day F_1,7_=0.7, p=0.44) and RNF (**H**, 15-min slot F_2,14_=10.7, p<0.01, despite of IVSA day × 15-min slot interaction F_1,10_=0.9, p=0.40, IVSA day F_1,7_=0.9, p=0.37) during the 1-hr IVSA sessions on both Day 1 and Day 5. **I-K**, Individual mouse values (small circles) and summarized data (large circles) from cocaine mice showing higher number of ALP (**I**, IVSA day F_1,9_=5.3, p=0.047, despite of IVSA day × 15-min slot interaction F_2,17_=1.3, p=0.34, 15-min slot F_2,18_=0.6, p=0.54) and RNF (**K**, IVSA day F_1,9_=5.5, p=0.04, despite of IVSA day × 15-min slot interaction F_3,27_=0.5, p=0.69, 15-min slot F_3,27_=0.2, p=0.90), but not ILP (**J**, IVSA day × 15-min slot interaction F_2,22_=1.0, p=0.39, IVSA day F_1,9_=1.7, p=0.22, 15-min slot F_2,16_=1.5, p=0.24), at the early time of the 1-hr IVSA sessions on Day 5, relative to that on Day 1. **L-Q,** Summarized data (**L-N**) and the fitted curves (**O-Q** by non-linear curve fitting in Prism 10, GraphPad) showing the significant interactions of three factors (i.e., saline/cocaine denoted “S/C”, IVSA days denoted “D”, and the minute during the 1-hr IVSA session denoted “min”) on ALP (**L**, S/C × D × min interaction F_119,1904_=1.5, p<0.01; S/C × D interaction F_1,16_=2.1, p=0.16; S/C × min interaction F_119,1904_=2.8, p<0.01; D × min interaction F_119,1904_=6.5, p<0.01; S/C F_1,16_=13.8, p<0.01; D F_1,16_=34.9, p<0.01; min F_119,1904_=224.5, p<0.01) and RNF (**M**, S/C × D × min interaction F_119,1904_=2.2, p<0.01; S/C × D interaction F_1,16_=5.3, p=0.03; S/C × min interaction F_119,1904_=3.0, p<0.01; D × min interaction F_119,1904_=9.6, p<0.01; S/C F_1,16_=15.8, p<0.01; D F_1,16_=44.1, p<0.01; min F_119,1904_=297.0, p<0.01), but not ILP (**N**, S/C × D × min interaction F_119,1904_=0.9, p=0.65; S/C × D interaction F_1,16_<0.01, p=0.93; S/C × min interaction F_119,1904_=1.5, p<0.01; D × min interaction F_119,1904_=4.0, p<0.01; S/C F_1,16_=2.3, p=0.15; D F_1,16_=2.7, p=0.12; min F_119,1904_=82.3, p<0.01). **R-U**, Scatter plots of the inter-ALP interval (y axis, in which 1000 sec was the max interval to the cutoff interval included) during the first (left in each panel) or the last (right in each panel) 15 min in the 1-hr IVSA training (x axis), together with the lines obtained by simply linear regression. In saline mice, significant non-zero linear regression was detected in the first 15 min (left panel in **R**, y=24x − 25, F_1,25_=9.3, p<0.01; r=0.52, p<0.01) but not the last 15 min (right panel in **R**, y=11x − 150, F_1,4_=0.1, p=0.75; r=0.17, p=0.75) during the 1-hr IVSA session on Day 1, whereas neither first 15 min (left panel in **S**, y=3x + 35, F_1,47_=1.0, p=0.32; r=0.15, p=0.32) nor the last 15 min (right panel in **S**, no data points available in this time window) showed any non-zero linear correlations on Day 5 of the 5-day saline IVSA sessions. In cocaine mice, significant non-zero linear regression was detected in the first 15 min but not the last 15 min during the 1-hr IVSA session on both Day 1 (left panel in **T** for the first 15 min, y=21x − 13, F_1,39_=13.7, p<0.01; r=0.51, p<0.01; right panel in **T** for the last 15 min, y= −6x − 484, F_1,59_=1.5, p=0.22; r= −0.16, p=0.22) and Day 5 (left panel in **U** for the first 15 min, y=9x + 37, F_1,68_=9.7, p<0.01; r=0.35, p<0.01; right panel in **U** for the last 15 min, y=−2x + 209, F_1,57_=0.1, p=0.74; r=−0.04, p=0.74) of the 5-day cocaine IVSA sessions. Data were analyzed by unpaired Student’s t test (**B**), two-way ANOVA with repeated measures on (a) Days and ALP/ILP (**C, D**), (b) Days (**E**) or (c) Day1/5 and different 15-min slots (**F-K**), three-way ANOVA with repeated measures on D and min (**L-N**), and simply linear regression and Pearson correlations (**R-U**). n=8 for saline mice, and n=10 for cocaine mice. In **F-K**, *, p<0.05; **, p<0.01, compared to the first 15 min (i.e., 0-15) on the same day; ^#^, p<0.05; ^##^, p<0.01, compared at the same 15-min slot between Day 1 *vs*. Day 5 (i.e., 0-15). In **R-U**, *, p<0.05; **, p<0.01.

**Figure 2.**
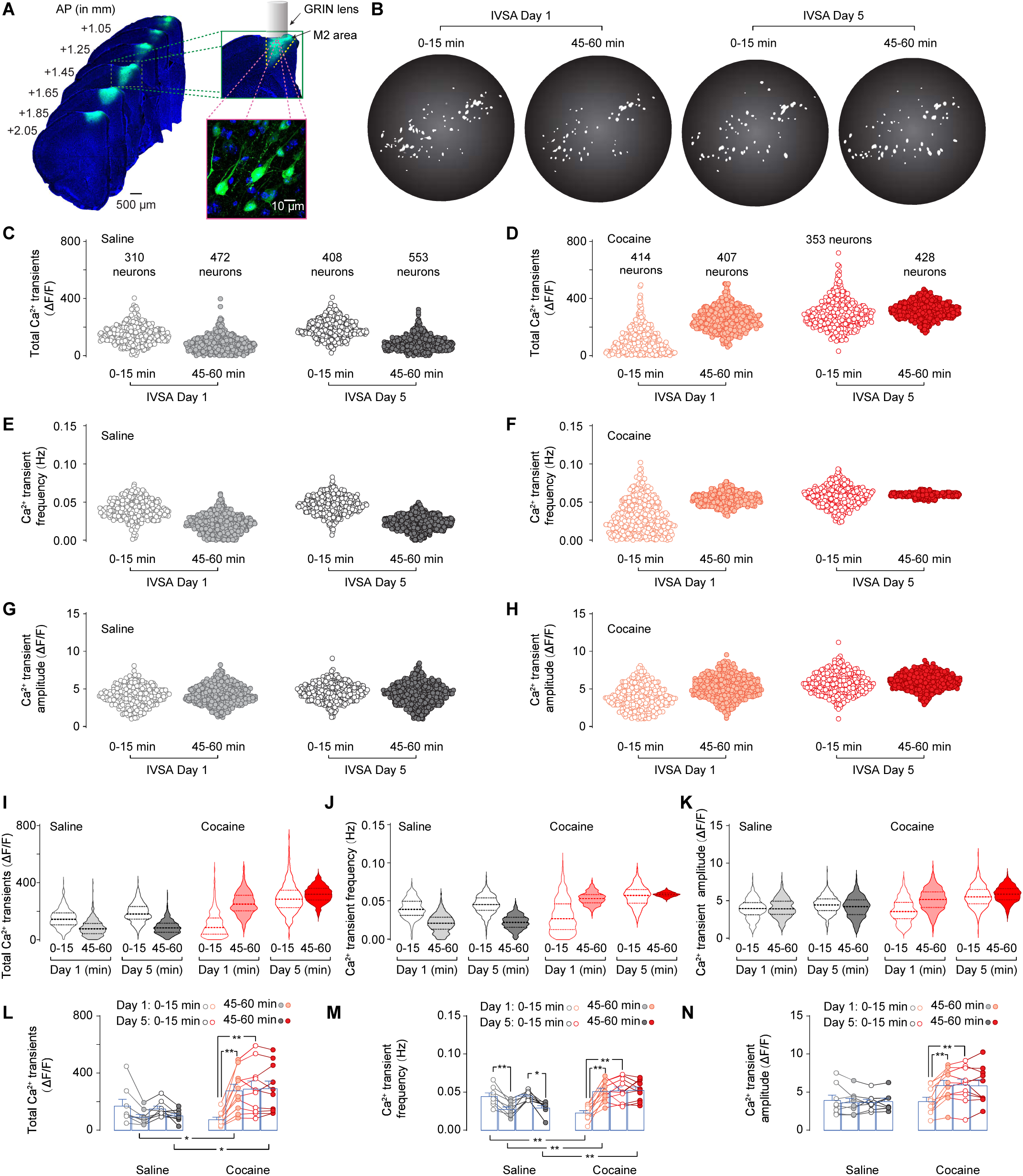
Ca^2+^ transients in M2 neurons in the first *vs*. last 15 min on Day 1 and Day 5 of IVSA. **A**, Representative images of GCaMP8f in M2 neurons, primarily the pyramidal projection neurons as demonstrated in the lower right panel. **B**, Images of maximum projections of *in vivo* Ca^2+^ images from a representative mouse. **C-H**, Scattered plots showing the total Ca^2+^ transients (**C**, **D**), and their frequency (**E**, **F**) and amplitude (**G**, **H**) in all the Ca^2+^ *in vivo* recording-captured M2 neurons in the first (open circles) and the last (solid circles) 15-min on the first and the last IVSA days in saline (**C**, **E**, **G**) and cocaine (**D**, **F**, **H**) mice. **I-K**, Violin plots demonstrating the density of detected M2 neurons at different levels of total Ca^2+^ transients (**I**), and their frequency (**J**) and amplitude (**K**) in the first (open plot) and the last (solid plot) 15-min on the first and the last IVSA days in saline (in grayish) and cocaine (in reddish) mice. There are three dashed lines splitting each violin plot at the quartile positions. **L-N**, Total Ca^2+^ transient (**L**), and their frequency (**M**) and amplitude (**N**) for each animal in the first (open plot) and the last (solid plot) 15-min on the first and the last IVSA days in saline (in grayish) and cocaine (in reddish) mice. The # of neurons are provided in **C,D**. Data from each neuron in **C-K** or from each animal in **L-N** were analyzed by three-way ANOVA, followed by Bonferroni’s post hoc test as shown in **Table S1** (**C-K**) and **S2** (**L-N**). In **L-N**, *, p<0.05; **, p<0.01.

## RESULTS

### IVSA events (i.e., ALP, ILP, RNF) during the IVSA sessions

Mice with pre-implanted GRIN lenses (n=18) into M2 and catheter into the jugular vein were randomly assigned to IVSA of saline or cocaine (**Fig. 1A**). Mice in these two groups demonstrated similar distance traveled during the 5-min open-field test (**Fig. 1B**), indicating their general locomotion was comparable. As expected, saline mice showed similar #s of ALP and ILP (**Fig. 1C**), whereas cocaine mice performed more on ALP, relative to ILP throughout the five 1-hr daily sessions (**Fig. 1D**). Thus, compared to saline mice, cocaine mice earned a high # of IV injections (i.e., RNFs; **Fig. 1E**). In order to evaluate the behavioral adaptations within one daily session, we divided the 1-hr daily session into 4 blocks of 15 mins each. It was apparent that, different from the gradual reduction of ALP, ILP and RNF from the first to the last 15-min block on both Day 1 and Day 5 of IVSA in saline mice (**Fig. 1F-H**), cocaine mice consistently demonstrated active ALP and RNF, accompanied with a low number of ILP, throughout the four 15-min blocks on both Day 1 and Day 5 of IVSA (**Fig. 1I-N**). The cumulative probability of three featured IVSA events (**i.e., ALP, ILP, RNF** in **Figs. 1L-Q**) revealed distinguishable time-courses of ALP and RNF, but not ILP, between saline *vs*. cocaine mice. Then the inter-ALP-interval (IAI; cut off at 1000 sec) was used as an example IVSA event to extract more details of the time courses of IVSA behaviors (**Fig. 1R-U**). Specifically, saline mice exhibited a linear increase in IAI during the first 15 min on Day 1, but not on Day 5, with very few or no within-1000-sec IAI occurrences in the last 15 min on both days. In contrast, cocaine mice showed a linear increase in IAI during the first 15 min on both Day 1 and Day 5. Although no linear changes in IAI were detected in the last 15 min, the number of ALP events remained constant during the last 15 min on both IVSA days. Our detailed analysis of mouse IVSA behaviors highlighted differences in cocaine taking within the 1-hr daily session (i.e., the first *vs*. the last 15 min) and across days (e.g., Day 1 *vs*. Day 5). To investigate the underlying neuronal alterations associated with these behaviors, we examined *in vivo* Ca^2+^ transients in M2 neurons both within a 1-hr daily session and across days, as detailed below.

### Alterations of the Ca^2+^ transients in M2 during IVSA in saline and cocaine mice

The GCaMP8f-expressing M2 neurons (**Fig. 2A, B**) appear predominantly the pyramidal projection neurons based on our observation that no less than 95% of these GCaMP8f-positive neurons detected in brain slices exhibited futures, including (1) pyramid-like soma, (2) apical dendrite from the apex of the cell body and extending toward the cortical surface, (3) basal dendrites from the base of the cell body, and (4) the typical dendritic arborization. The activity of these GCaMP8f-expressing neurons, particularly within Layer 2, a layer we recently identified as critical in addiction-like behaviors (Huang and Ma, 2023a), was monitored in real time during IVSA. We quantified the total Ca^2+^ transients within a specific time window by summing the amplitude of all observed Ca^2+^ transients in that time window. Additionally, we calculated the frequency in Hz, and the average amplitude in that time window by dividing the total Ca^2+^ transients (i.e., the sum of the amplitude of all Ca^2+^ transients in the time window) by the number of Ca^2+^ transients. Changes of Ca^2+^ transients, particularly their frequency, in saline *vs*. cocaine mice were observed when neuron was used as the statistical unit (**Fig. 2C-H**), with expected higher statistical significance (**Table S1**), relative to data analyzed as animal-based unit (**Fig. 2L-N**), due to a larger number of neurons (i.e., 310 - 553) per experimental condition. Specifically, using animal as the statistical unit, we found decreases in the total Ca^2+^ transients, which were primarily attributable to reduction of the frequency, but not amplitude, of Ca^2+^ transients from the first 15-min to the last 15-min during the 1-hr IVSA session on both Day 1 and Day 5 in saline mice. In contrast, there were increases of total Ca^2+^ transients in cocaine mice, which were attributable to both frequency and amplitude of the Ca^2+^ transients on Day 1, and then remained at high levels consistently during the 1-hr IVSA session on Day 5. The reduction of the non-reinforcing operant behavior (i.e., ALP and ILP in saline mice as shown in **Fig. 1F, G**) from the early to the late stage of daily session may be associated with a reduction in the frequency of Ca^2+^ transients. This reduction of Ca^2+^ transients in saline mice could be the result of M2 neuronal adaptations to the early non-reinforced operant behaviors and also the cause of less operant behaviors in the late stage of the daily sessions. On the contrary, the reinforced operant behavior (i.e., IV cocaine-associated ALP in cocaine mice) was accompanied by lower Ca^2+^ transient frequency in M2 neurons in the early stage (i.e., the first 15 min of the 1-hr IVSA session) on Day 1, with a significant rebound in the late stage (i.e., the last 15 min of the 1-hr IVSA session) on Day 1 and persisted at a high level throughout the 1-hr IVSA session on Day 5. This pattern suggests different M2 neuron engagement in the early *vs*. late phase of cocaine IVSA, potentially corresponding to the transition from drug use to drug misuse.

In order to determine the associations between an IVSA event (i.e., ALP, ILP, or RNF) and M2 Ca^2+^ transients (including the total transients, frequency, and amplitude), analyses of the Ca^2+^ transients within the 3 min before ALP/ILP or after RNF were performed by averaging the IVSA event-based 3-min Ca^2+^ transients within the first and the last 15 min of the 1-hr IVSA session on Day 1 and Day 5 in Figs 3-5, S1-S6 as described in the following two subsections.

**Figure 3.**
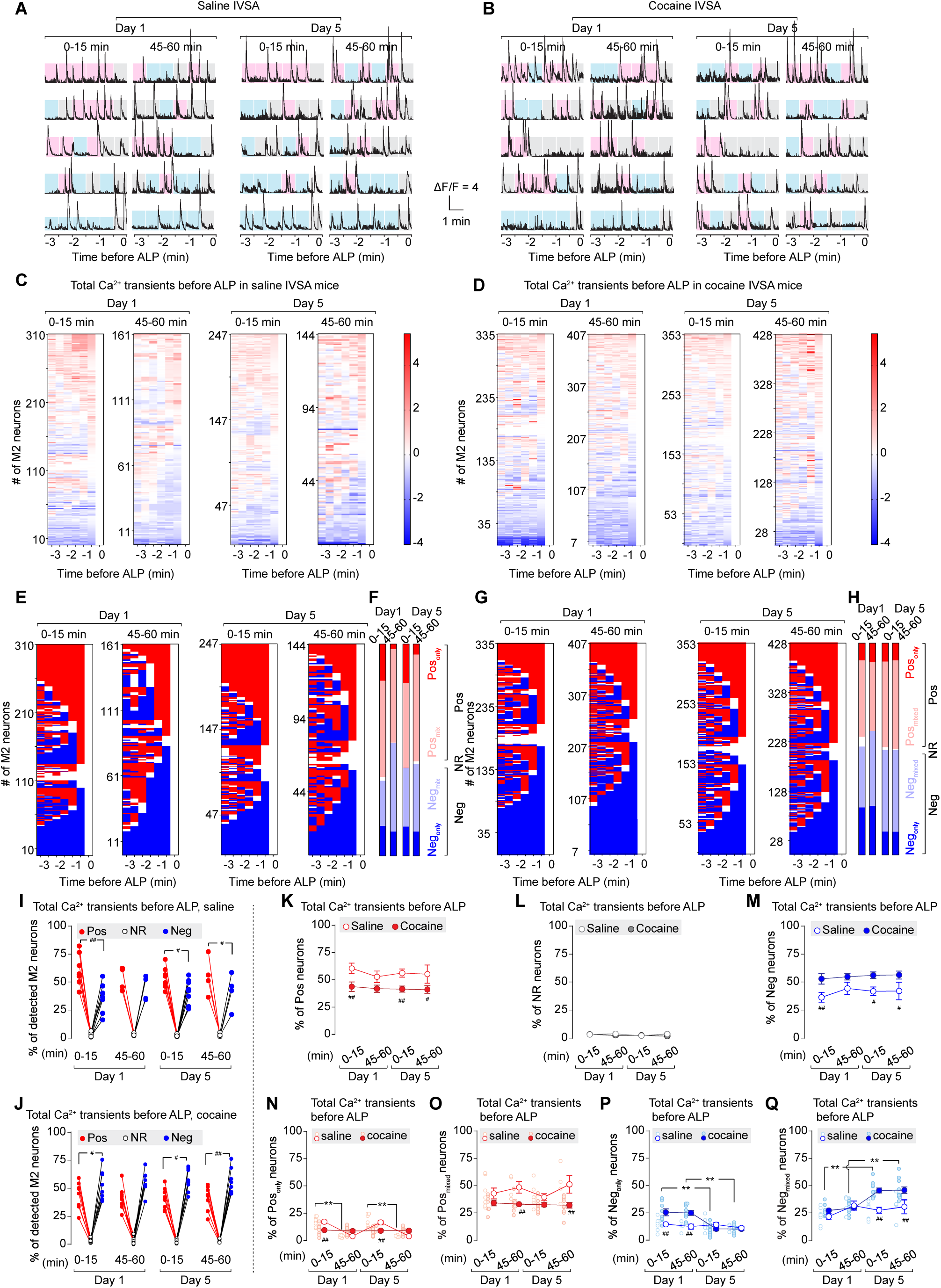
Ca^2+^ transients in M2 neurons during 3 min before ALP in saline vs. cocaine mice. **A**, **B**, Representative Ca^2+^ transient traces within 3 min before ALP in the first (left in each pair of the trace columns) and the last (right in each pair of the trace columns) 15 min during Day 1 and Day 5 IVSA sessions in saline (**A**) and cocaine (**B**) mice. Each of the 30-sec blocks are shadowed in gray, pink, or blue, indicating no significant difference, positive response, or negative response, respectively, as determined by the ΔZ score of that 30-sec block. **C, D**, Heat maps of ΔZ scores of neuronal total Ca^2+^ transients in detected M2 neurons displayed by rows of six 30-sec blocks within 3 min before ALP in the first (left in each pair of the trace columns) and the last (right in each pair of the trace columns) 15 min during Day 1 and Day 5 IVSA sessions in saline (**C**) and cocaine (**D**) mice. **E-H**, Total Ca^2+^ transient ΔZ based-classification of **each** 30-sec block of M2 neurons within 3 min before ALP in the first (left in each pair of the trace columns) *vs*. the last (right in each pair of the trace columns) 15 min during Day 1 and Day 5 IVSA sessions in saline (**E**) and cocaine (**G**) mice. The proportions of M2 neurons in each category in a 15-min session are summarized in **F** for saline mice and **H** for cocaine mice. **I**, **J**, Summarized data showing the percentage of three ALP & total Ca^2+^ transient - based categories of M2 neurons in saline (**I**) and cocaine (**J**) mice. Both saline (**I**, neuron type × IVSA training session interaction F_6,40_=0.4, p=0.86; neuron type F_1,20_=103.2, p<0.01; and IVSA training session F_3,20_=0.0, p>0.99) and cocaine (**J**, neuron type × IVSA training session interaction F_6,66_=0.2, p=0.98; neuron type F_1,35_=238.0, p<0.01; and IVSA training session F_3,33_=0.9, p=0.48) mice demonstrated different proportions among three neuronal types. **K-M**, Summarized data showing smaller proportions of ALP & total Ca^2+^ transient - categorized Pos neurons (**K**, saline/cocaine × IVSA training session interaction F_3,53_=0.1, p=0.93; saline/cocaine F_1,53_=21.0, p<0.01; and IVSA training session F_3,53_=0.5, p=0.69), similar proportions of ALP-categorized NR neurons (**L**, saline/cocaine × IVSA training session interaction F_3,53_=0.1, p=0.98; saline/cocaine F_1,53_=0.2, p=0.67; and IVSA training session F_3,53_=0.9, p=0.46), and larger proportions of ALP-categorized Neg neurons (**M**, saline/cocaine × IVSA training session interaction F_3,53_=0.2, p=0.93; saline/cocaine F_1,53_=20.0, p<0.01; and IVSA training session F_3,53_=0.6, p=0.61) in cocaine mice, relative to that in saline mice. **N**, **O**, Summarized data showing training session-associated differences in the proportions of ALP & total Ca^2+^ transient -categorized Pos_only_ neurons (**N**, saline/cocaine × IVSA training session interaction F_3,53_=11.7, p<0.01; saline/cocaine F_1,53_=1.7, p=0.19; and IVSA training session F_3,53_=13.2, p<0.01), and ALP-categorized Pos_mixed_ neurons (**O**, saline/cocaine × IVSA training session interaction F_3,53_=1.3, p=0.29; saline/cocaine F_1,53_=24.8, p<0.01; and IVSA training session F_3,53_=1.1, p=0.36) between saline *vs*. cocaine mice. **P**, **Q**, Summarized data showing training session-associated differences in the proportions of ALP & total Ca^2+^ transient -categorized Neg_only_ neurons (**P**, saline/cocaine × IVSA training session interaction F_3,53_=7.0, p<0.01; saline/cocaine F_1,53_=8.5, p<0.01; and IVSA training session F_3,53_=8.5, p<0.01), and ALP-categorized Neg_mixed_ neurons (**Q**, saline/cocaine × IVSA training session interaction F_3,53_=4.5, p=0.01; saline/cocaine F_1,53_=18.0, p<0.01; and IVSA training session F_3,53_=8.9, p<0.01) between saline *vs*. cocaine mice. Data were analyzed by two-way ANOVA with repeated measures on neuron types and IVSA sessions (**I**, **J**), on IVSA sessions (**K-Q**), followed by Bonferroni *post hoc* test. Between different IVSA sessions, *, p<0.05; **, p<0.01. Between saline vs. cocaine, #, p<0.05; ##, 0.01.

### Cocaine affected Ca^2+^ transients before ALP, but not ILP

As detailed in **MATERIALS AND METHODS**, the 3-min window of M2 neurons associated with ALP or ILP, which means the 6^th^ 30-sec block (i.e., −30 to 0 sec or −0.5 to 0 min) was the block when the ALP or ILP occurred, was categorized as (a) positive (Pos), (b) non-responsive (NR), and (c) negative (Neg) if the 5^th^ 30-sec block (i.e. −60 to - 30 sec or −1 to 0.5 min) was classified as positive, non-responsive, and negative, respectively. M2 neuronal categorization based on the total Ca^2+^ transients within-3 min before ALP/ILP disclosed a lower Pos% (41-44% as a range of the mean values; the same below) and higher Neg% (53-56%) before ALP in cocaine mice, relative to that in saline mice (Pos, 53-60%; Neg, 36-44%) (**Fig. 3A-M**), but no significant differences of Pos% (saline, 52-56%; cocaine, 43-52%) or Neg% (saline, 44-46%; cocaine, 48-56%) before ILP between saline *vs*. cocaine mice (**Fig.4A-M**). Further analysis of the subtypes of Pos and Neg neurons in M2 revealed that the decrease in Pos% before ALP in cocaine mice was attributable to lower Pos_only_% in the first 15 min and lower Pos_mixed_% in the last 15 min on both Day 1 and Day 5 (**Fig. 3N, O**). The increases of Neg% before ALP in cocaine mice were attributable to higher Neg_only_% but not Neg_mixed_% on Day 1 and higher Neg_mixed_% but not Neg_only_% on Day 5 (**Fig. 3P, Q**). Interestingly, the within-session fluctuation of Pos_only_% and Pos_mixed_% was observed before ALP but not ILP in saline mice and before ILP but not ALP in cocaine mice (**Figs. 3N, O; 4N, O**). No within-session fluctuations of Neg subtype neurons (Neg_only_ and Neg_mixed_) were observed in either saline or cocaine mice before ALP and ILP (**Figs. 3P, Q; 4P, Q**). When we investigated the Ca^2+^ frequency and the amplitude-based M2 neuronal categorization, differences of the M2 neuronal categorization before ALP/ILP between saline *vs*. cocaine mice should be attributed to both frequency (**Figs S1, S3**) and amplitude (**Figs S2, S4**). In summary, we **conclude** that the sub-population categories of M2 neurons are distinguishable between saline *vs*. cocaine mice primarily before ALP but not before ILP. In saline mice, we observed the opposite fluctuations of the ALP-associated neuronal ratio (1) between Pos_only_ *vs*. Pos_mixed_ subtype neurons within the 1-hr daily session on both days, and (2) between-day, but not within-session, alterations in the ratio between Neg_only_ vs. Neg_mixed_ subtype neurons. This leads to our **second conclusion** that cocaine IVSA abrogated the within-session and between-day adaptations before ALP observed in saline mice, resulting in a flat ratio curve of all the four subtypes of M2 neurons throughout the IVSA training sessions. The consistent cocaine taking behaviors during the 1-hr daily sessions on both Day 1 and Day 5 might be attributable to this flat ratio curve.

**Figure 4.**
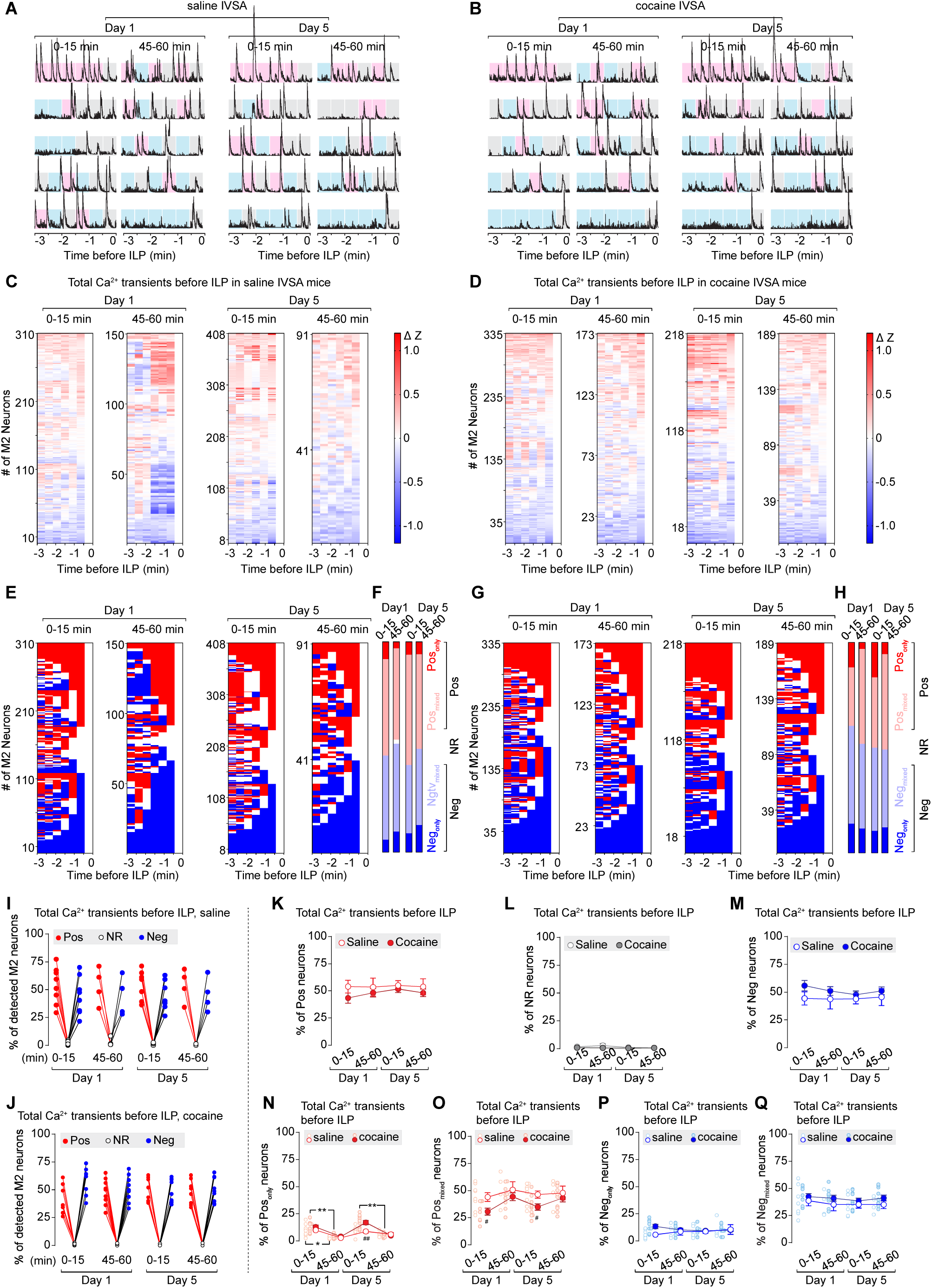
Ca^2+^ transients in M2 neurons during the 3 min before ILP in saline vs. cocaine mice. **A**, **B**, Representative Ca^2+^ transient traces within 3 min before ILP in the first (left in each pair of the trace columns) and the last (right in each pair of the trace columns) 15 min during Day 1 and Day 5 IVSA sessions in saline (**A**) and cocaine (**B**) mice. Each of the 30-sec blocks are shadowed in gray, pink, or blue, indicating the no significant difference, positive response, or negative response, respectively, as sorted by the ΔZ score of that 30-sec block. **C, D**, Heat maps of ΔZ scores of neuronal total Ca^2+^ transients in detected M2 neurons displayed by rows of six 30-sec blocks within 3 min before ILP in the first (left in each pair of the trace columns) and the last (right in each pair of the trace columns) 15 min during Day 1 and Day 5 IVSA sessions in saline (**C**) and cocaine (**D**) mice. **E-H**, Total Ca^2+^ transient ΔZ-based classification of **each** 30-sec block of M2 neurons within 3 min before ILP in the first (left in each pair of the trace columns) *vs*. the last (right in each pair of the trace columns) 15 min during Day 1 and Day 5 IVSA sessions in saline (**E**) and cocaine (**G**) mice. The proportions of M2 neurons in each category in a 15-min session is summarized in **F** for saline mice and **H** for cocaine mice. **I**, **J**, Summarized data showing the percentage of three ILP & total Ca^2+^ transient - based categories of M2 neurons in saline (**I**) and cocaine (**J**) mice. Both saline (**I**, neuron type × IVSA training session interaction F_6,40_<0.1, p>0.99; neuron type F_1,20_=71.8, p<0.01; and IVSA training session F_3,20_=155.4, p<0.01) and cocaine (**J**, neuron type × IVSA training session interaction F_6,66_=0.2, p=0.98; neuron type F_1,33_=222.0, p<0.01; and IVSA training session F_3,33_=0.5, p=0.69) mice demonstrated different proportions among three neuronal types. **K-M**, Summarized data showing similar proportions of ILP & total Ca^2+^ transient - categorized Pos neurons (**K**, saline/cocaine × IVSA training session interaction F_3,53_=0.2, p=0.87; saline/cocaine F_1,53_=3.1, p=0.08; and IVSA training session F_3,53_=0.4, p=0.77), ILP-categorized NR neurons (**L**, saline/cocaine × IVSA training session interaction F_3,53_=1.2, p=0.31; saline/cocaine F_1,53_=3.2, p=0.08; and IVSA training session F_3,53_=1.6, p=0.19), and ILP-categorized Neg neurons (**M**, saline/cocaine × IVSA training session interaction F_3,53_=0.2, p=0.86; saline/cocaine F_1,53_=3.7, p=0.06; and IVSA training session F_3,53_=0.3, p=0.82) between saline *vs*. cocaine mice. **N**, **O**, Summarized data showing significant differences in the proportions of ILP & total Ca^2+^ transient -categorized Pos_only_ neurons (**N**, saline/cocaine × IVSA training session interaction F_3,53_=3.4, p=0.03; saline/cocaine F_1,53_=6.0, p=0.02; and IVSA training session F_3,53_=16.0, p<0.01), and ILP-categorized Pos_mixed_ neurons (**O**, saline/cocaine × IVSA training session interaction F_3,53_=0.5, p=0.68; saline/cocaine F_1,53_=9.3, p<0.01; and IVSA training session F_3,53_=2.4, p=0.08) between saline *vs*. cocaine mice. **P**, **Q**, Summarized data showing no detectable differences in the proportions of ILP & total Ca^2+^ transient-categorized Neg_only_ neurons (**P**, saline/cocaine × IVSA training session interaction F_3,53_=2.0, p=0.12; saline/cocaine F_1,53_=2.7, p=0.11; and IVSA training session F_3,53_=0.2, p=0.93), or ILP-categorized Neg_mixed_ neurons (**Q**, saline/cocaine × IVSA training session interaction F_3,53_=0.1, p=0.97; saline/cocaine F_1,53_=3.6, p=0.06; and IVSA training session F_3,53_=0.5, p=0.71) between saline *vs*. cocaine mice. Data were analyzed by two-way ANOVA with repeated measures on neuron types and IVSA sessions (**I**, **J**), on IVSA sessions (**K-Q**), followed by Bonferroni *post hoc* test. Between different IVSA sessions, *, p<0.05; **, p<0.01. Between saline vs. cocaine, #, p<0.05; ##, 0.01.

### Cocaine IVSA affected Ca^2+^ transients after RNF

As detailed in **MATERIALS AND METHODS**, the 3-min window after RNF, which means the 1^st^ 30-sec block (i.e., 0 to 30 sec or 0 to 0.5 min) was the block when the RNF occurred, was categorized as positive (Pos) and negative (Neg) if the nearest responsive 30-sec block was classified as positive and negative, respectively (**Fig. 5A-M**). The 3-min window associated with RNF was categorized as not responsive (NR) if all five 30-sec blocks from 30 to 180 sec (i.e., 0.5 to 3 min) were classified as non-responsive. M2 neuronal categorization based on the total Ca^2+^ transients within the 3-min after RNF revealed that the % of Pos, NR and Neg neurons in M2 remained consistent across the 1-hr daily sessions on both Day 1 and Day 5 of saline IVSA, although the Ca^2+^ transients after RNF showed a higher % of Pos neurons (i.e., 59-64%) compared to the Neg neurons (i.e., 29-34%). M2 neurons in cocaine mice demonstrated lower Pos% (i.e., 32%) in the first 15 min which was then increased in the last 15 min (i.e., 49% as mean) on Day 1, and stayed at high levels in the first (58%) and the last (60%) 15 min on Day 5. On the contrary, Neg% in cocaine mice was detected with reductions on Day 1 (from the first to the last 15 min: 58% to 47%), followed by a consistent low level on Day 5 (from the first to the last 15 min: 37% to 36 %). Further dissection of the subtypes of Pos and Neg neurons in M2 in cocaine mice demonstrated that the increases of Pos% were attributable to the changes in Pos_mixed_% but not Pos_only_%, and decreases of Neg% were attributable to the alteration of Neg_only_% but not Neg_mixed_% (**Fig. 5N-Q**). Relative to that in saline mice, the lower Pos_mixed_% and higher Neg_only_% in the first 15 min on Day 1 after cocaine IV injection may indicate the sensitivity of cocaine mice to the IV cocaine delivery at the early stage, which was altered to the level closer to that in saline mice in the last 15 min on Day 1, and then no more differences between saline *vs*. cocaine mice occurred throughout the 1-hr IVSA session on Day 5. Different from what we observed in the M2 neuronal categorization before ALP/ILP, the differences of M2 neuronal categorization after RNF between saline *vs*. cocaine mice should be attributed to frequency (**Figs S5**) but not amplitude (**Figs S6**). The cocaine-IV associated higher Neg_only_% and lower Pos_mix_% at the early IVSA stage indicated temporary inhibition of IV cocaine on Ca^2+^ transients in M2.

**Figure 5.**
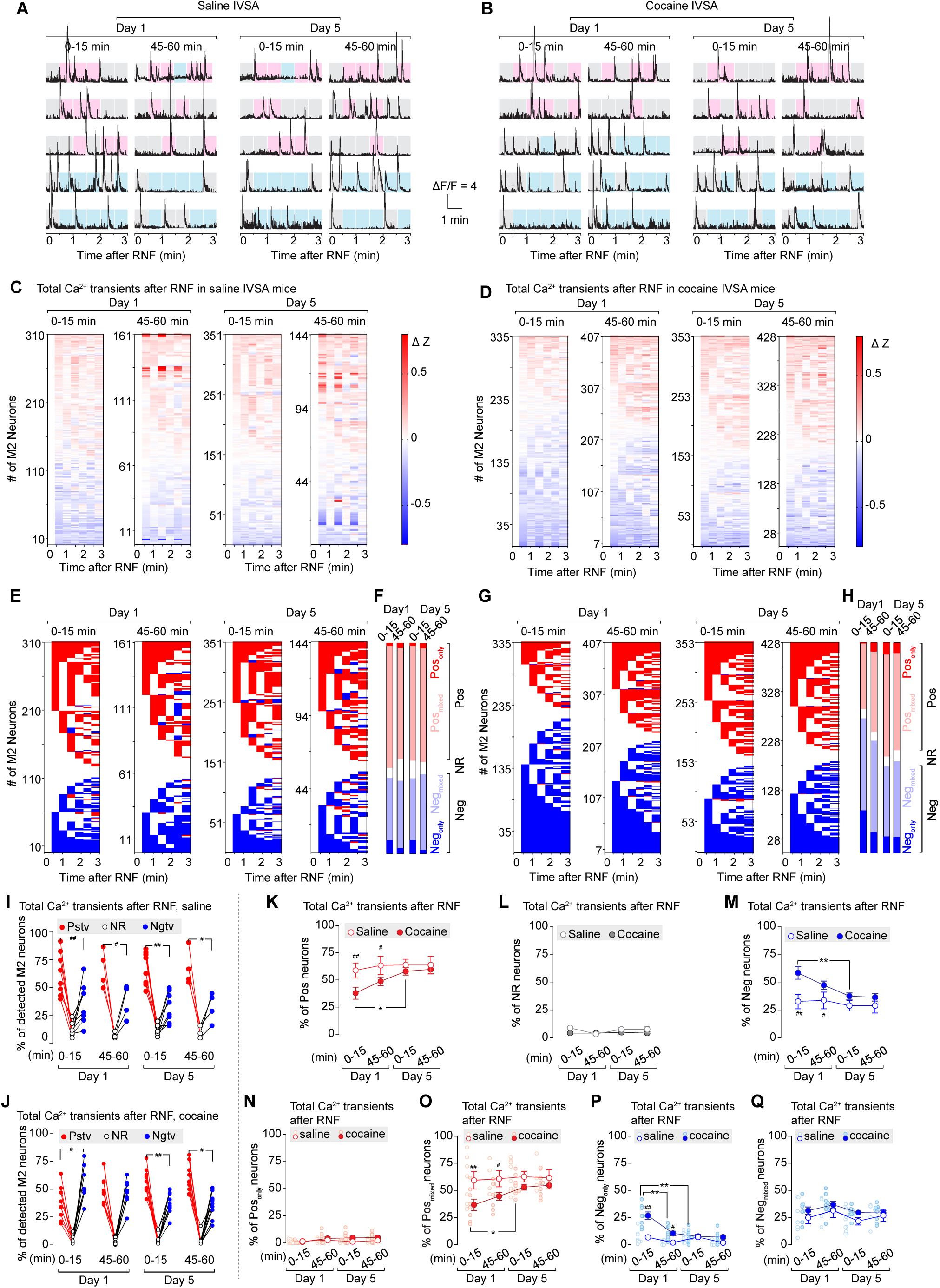
Ca^2+^ transients in M2 neurons during the 3 min after RNF in saline vs. cocaine mice. **A**, **B**, Representative Ca^2+^ transient traces within 3 min after RNF in the first (left in each pair of the trace columns) and the last (right in each pair of the trace columns) 15 min during Day 1 and Day 5 IVSA sessions in saline (**A**) and cocaine (**B**) mice. Each of the 30-sec blocks is shadowed in gray, pink, or blue, indicating the no significant difference, positive response, or negative response, respectively, as sorted by the ΔZ score of that 30-sec block. **C, D**, Heat maps of ΔZ scores of neuronal total Ca^2+^ transients in detected M2 neurons displayed by rows of six 30-sec blocks within 3 min after RNF in the first (left in each pair of the trace columns) and the last (right in each pair of the trace columns) 15 min during Day 1 and Day 5 IVSA sessions in saline (**C**) and cocaine (**D**) mice. **E-H**, Total Ca^2+^ transient ΔZ-based classification of **each** 30-sec block of M2 neuron within 3 min after RNF in the first (left in each pair of the trace columns) *vs*. the last (right in each pair of the trace columns) 15 min during Day 1 and Day 5 IVSA sessions in saline (**E**) and cocaine (**G**) mice. The proportions of M2 neurons in each category in a 15-min session are summarized in **F** for saline mice and **H** for cocaine mice. **I**, **J**, Summarized data showing the percentage of three RNF & total Ca^2+^ transient - based categories of M2 neurons in saline (**I**) and cocaine (**J**) mice. Both saline (**I**, neuron type × IVSA training session interaction F_6,40_=0.2, p=0.98; neuron type F_1,23_=59.2, p<0.01; and IVSA training session F_3,20_<0.1, p>0.99) and cocaine (**J**, neuron type × IVSA training session interaction F_6,66_=5.7, p<0.01; neuron type F_1,39_=149.3, p<0.01; and IVSA training session F_3,33_=128.9, p<0.01) mice demonstrated different proportions among three neuronal types. **K-M**, Summarized data showing lower proportions of RNF & total Ca^2+^ transient - categorized Pos neurons (**K**, saline/cocaine × IVSA training session interaction F_3,53_=1.1, p=0.36; saline/cocaine F_1,53_=8.4, p=0.01; and IVSA training session F_3,53_=2.8, p=0.049), similar proportions of RNF & total Ca^2+^ transient-categorized NR neurons (**L**, saline/cocaine × IVSA training session interaction F_3,53_=0.8, p=0.49; saline/cocaine F_1,53_=3.3, p=0.08; and IVSA training session F_3,53_=0.8, p=0.47), and higher proportions of RNF & total Ca^2+^ transient -categorized Neg neurons (**M**, saline/cocaine × IVSA training session interaction F_3,53_=1.5, p=0.22; saline/cocaine F_1,53_=14.7, p<0.01; and IVSA training session F_3,53_=3.2, p=0.03) in cocaine mice, relative to that in saline mice. **N**, **O**, Summarized data showing training session-associated differences in the proportions of RNF & total Ca^2+^ transient-categorized Pos_only_ neurons (**N**, saline/cocaine × IVSA training session interaction F_3,53_=1.3, p=0.29; saline/cocaine F_1,53_=4.4, p=0.04; and IVSA training session F_3,53_=1.6, p=0.21), and RNF-categorized Pos_mixed_ neurons (**O**, saline/cocaine × IVSA training session interaction F_3,53_=1.0, p=0.42; saline/cocaine F_1,53_=13.8, p<0.01; and IVSA training session F_3,53_=1.8, p=0.15) between saline *vs*. cocaine mice. **P**, **Q**, Summarized data showing training session-associated differences in the proportions of RNF & total Ca^2+^ transient -categorized Neg_only_ neurons (**P**, saline/cocaine × IVSA training session interaction F_3,53_=7.9, p<0.01; saline/cocaine F_1,53_=28.1, p<0.01; and IVSA training session F_3,53_=12.6, p<0.01), and RNF-categorized Neg_mixed_ neurons (**Q**, saline/cocaine × IVSA training session interaction F_3,53_=0.2, p=0.89; saline/cocaine F_1,53_=4.4, p=0.04; and IVSA training session F_3,53_=1.8, p=0.161) between saline *vs*. cocaine mice. Data were analyzed by two-way ANOVA with repeated measures on neuron types and IVSA sessions (**I**, **J**), on IVSA sessions (**K-Q**), followed by Bonferroni’s *post hoc* test. Between different IVSA sessions, *, p<0.05; **, p<0.01. Between saline vs. cocaine, #, p<0.05; ##, 0.01.

### Cumulative effects of IVSA events on Ca^2+^ transient activity

We would like to clarify that the Ca^2+^ transients within 3 min before ALP/ILP or after RNF-based neuronal categorization (**Figs. 3-5, S1-S6**) was determined using the ΔZ scores of Ca^2+^ transients in six 30-sec blocks, averaged from all of the IVSA events (i.e., ALP, ILP, RNF) during a 15-min time window. Thus, the sequence of the IVSA events was not considered in that data processing. Given that drug-taking behavior is a typical experience-dependent output, we proposed that the previous drug taking and drug delivery affected the upcoming neuronal activity involved in regulating impending drug taking behavior. In order to test whether M2 neurons were involved in mediating the cumulative effects of drug taking and drug delivery, we conducted linear regression analyses using two different approaches for the # of ALP, ILP and RNF events: (1) the counts exclusively within each 30-sec block, and (2) the counts within the current 30-sec block along with any previous 30-sec blocks, if applicable. Compared to the linear analysis using non-cumulative (nonCML) number of IVSA events, we detected a larger proportion of linearly associated neurons (LANs) in both saline and cocaine mice, particularly in the first 15 min on both Day 1 and Day 5, when the cumulative (CML) number of ALP (**Fig. 6A, B**) and RNF was used (**Fig. 6I, J**). Furthermore, the increases in LAN% associated with the CML numbers of ALP and RNF were significantly higher in cocaine mice than in saline mice (**Figs. 6C, D, K, L; S7**), indicating that cocaine taking behaviors can further enhance the sequence effects of ALP and RNF. The CML number of ILP-associated increases of the LAN% was only detected in saline mice but not cocaine mice (**Fig. 6E, F, H**). Thus, we conclude there is a significant sequence effect of ALP and RNF, which is more pronounced in cocaine mice. We then focused on the LANs identified by the CML number of IVSA events to address the following two questions.

**Figure 6.**
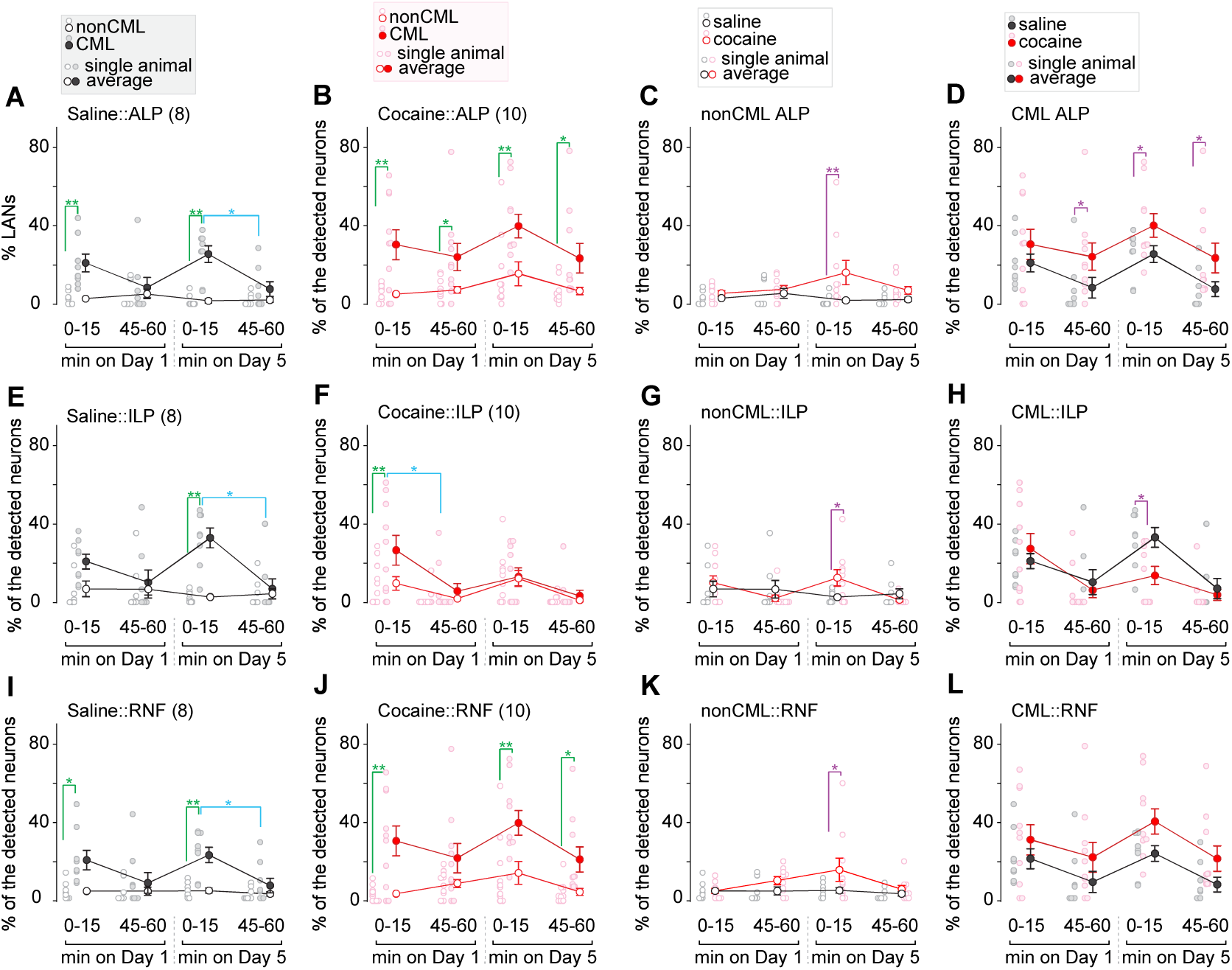
Non-cumulative *vs*. cumulative effects of cocaine-associated events on Ca^2+^ transients in M2 neurons. **A-D**, % of linearly associated neurons (LANs) analyzed with the non-cumulative (nonCML) and cumulative (CML) ALP #s on Ca^2+^ transients detected in M2 neurons. CML-LANs in M2 were detected with a higher number in both saline (**A**, nonCML/CML × IVSA training session interaction F_2,13_=3.9, p=0.047; nonCML/CML F_1,7_=44.8, p<0.01; and IVSA training session F_3,19_=4.0, p=0.03) and cocaine (**B**, nonCML/CML × IVSA training session interaction F_2,17_=2.3, p=0.13; nonCML/CML F_1,9_=21.0, p<0.01; and IVSA training session F_3,23_=0.5, p=0.67) mice. The % of nonCML LANs (**C**, saline/cocaine × IVSA training session interaction F_3,48_=1.6, p=0.20; saline/cocaine F_1,16_=16.2, p<0.01; and IVSA training session F_3,48_=0.9, p=0.44) and the CML-LANs (**D**, saline/cocaine × IVSA training session interaction F_3,48_=0.1, p=0.93; saline/cocaine F_1,16_=6.5, p=0.02; and IVSA training session F_3,48_=4.1, p=0.01) were higher in cocaine mice, relative to that in saline mice. **E-H**, % of LANs analyzed with nonCML and CML ILP #s on Ca^2+^ transients detected in M2 neurons. CML-LANs in M2 were detected with a higher number in both saline (**E**, nonCML/CML × IVSA training session interaction F_2,15_=2.2, p=0.14; nonCML/CML F_1,7_=19.4, p<0.01; and IVSA training session F_2,17_=6.3, p=0.01) and cocaine (**F**, nonCML/CML × IVSA training session interaction F_2,18_=6.3, p=0.01; nonCML/CML F_1,9_=6.7, p=0.03; and IVSA training session F_2,15_=2.0, p=0.17) mice. The % of nonCML LANs (**G**, saline/cocaine × IVSA training session interaction F_3,48_=2.1, p=0.12; saline/cocaine F_1,16_=0.5, p=0.48; and IVSA training session F_3,48_=1.3, p=0.27) and the CML LANs (**H**, saline/cocaine × IVSA training session interaction F_3,48_=2.2, p=0.10; saline/cocaine F_1,16_=1.7, p=0.21; and IVSA training session F_3,48_=7.5, p<0.01). **I-L**, % of LANs analyzed with the non-cumulative (nonCML) and cumulative (CML) RNF #s on Ca^2+^ transients detected in M2 neurons. CML-LANs in M2 were detected with a higher number in both saline (**I**, nonCML/CML × IVSA training session interaction F_2,16_=2.1, p=0.15; nonCML/CML F_1,7_=40.1, p<0.01; and IVSA training session F_2,13_=3.9, p=0.046) and cocaine (**J**, nonCML/CML × IVSA training session interaction F_2,19_=1.1, p=0.36; nonCML/CML F_1,9_=20.2, p<0.01; and IVSA training session F_2,16_=2.7, p=0.10) mice. The % of nonCML LANs (**K**, saline/cocaine × IVSA training session interaction F_3,48_=1.1, p=0.34; saline/cocaine F_1,16_=8.0, p=0.01; and IVSA training session F_3,48_=1.6, p=0.20) and the CML LANs (**L**, saline/cocaine × IVSA training session interaction F_3,48_=0.1, p=0.95; saline/cocaine F_1,16_=5.4, p=0.03; and IVSA training session F_3,48_=4.3, p=0.01) were higher in cocaine mice, relative to that in saline mice. Data were analyzed by two-way ANOVA with repeated measures, followed by Bonferroni’s post-test. *p<0.05, **p<0.01, marked in green as comparisons between nonCML *vs*. CML, blue as comparisons between different IVSA sessions, or purple as comparations between saline *vs*. cocaine.

**First**, was the increased LAN% in M2 positively or negatively associated between the Ca^2+^ influx and the IVSA event number? According to the Pearson correlation coefficient analyses, our results demonstrated that, unlike the reduced ratio of the LAN_Pos_% to LAN_Neg_% associated with ALP, ILP and RNFS from the first to the last 15 min on both Day 1 and Day 5 in saline mice, cocaine IVSA increased the ratio of the LAN_Pos_% to LAN_Neg_% of M2 neurons associated with the ALP and RNFS, but no significant changes in the ratio of LAN_Pos_% to LAN_Neg_% associated with ILP (**Figs. 7**). This increased ratio in cocaine mice was observed from the first to the last 15 min on Day 1 and then remained at a high level throughout the 1-hr session on Day 5.

**Figure 7.**
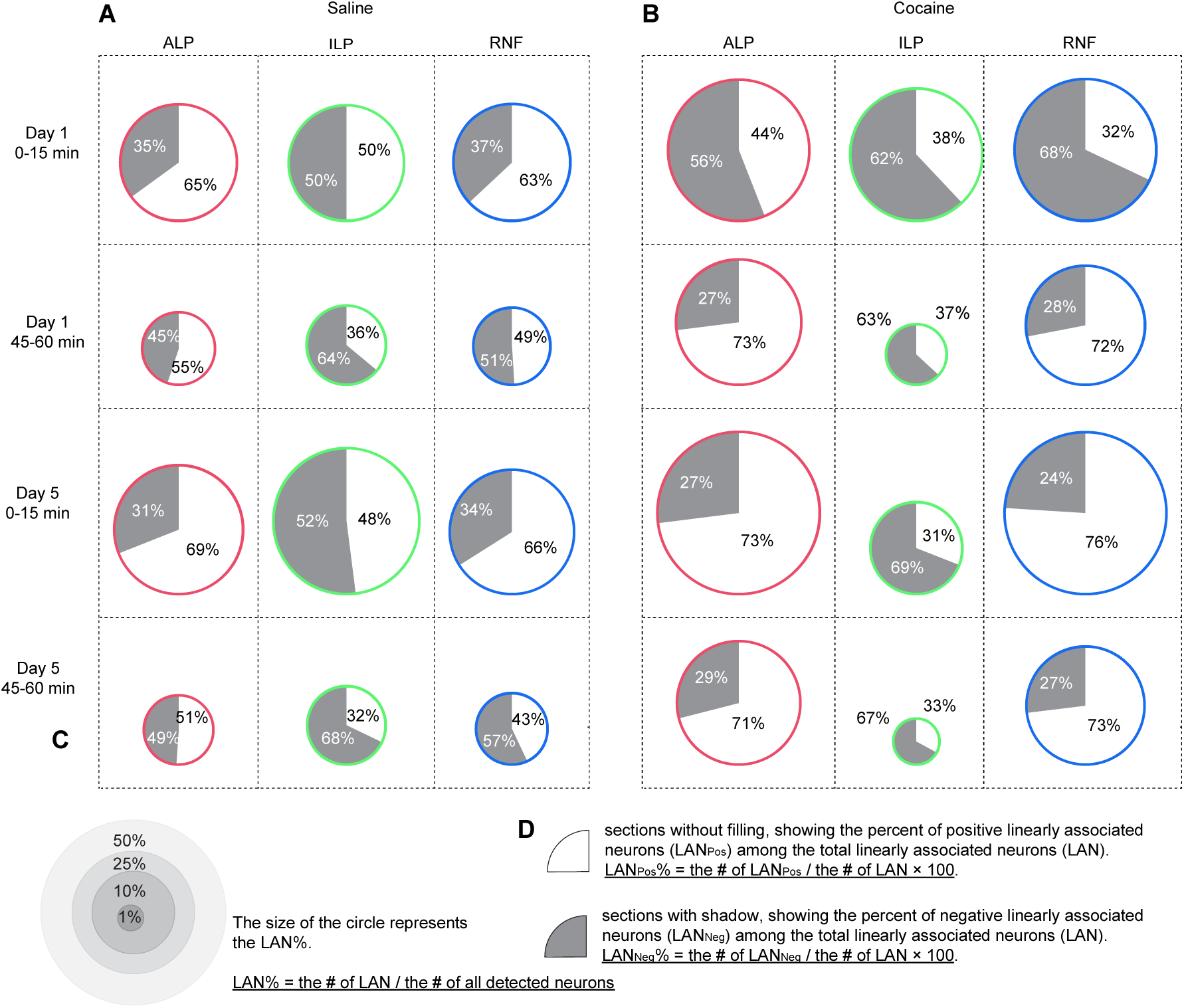
Percentage of positive *vs*. negative LANs in M2 during saline or cocaine taking. **A**, **B**, % of the positive (in sections with no filling) *vs*. negative (in sections with shadow) LANs identified by linear regression analyses of cumulative # of *in vivo* events, including ALP (left column), ILP (middle column) and RNF (right column), respectively, with the Ca^2+^ transient (i.e., the sum of the amplitude) in each 30-sec block from individual neurons in M2 during the first or the last 15 min on Day 1 or Day 5 of saline (**A**) or cocaine (**B**) IVSA. **C, D**, Figure legends demonstrating how to calculate (1) the % of LANs, which determines the size of each circle (**C**), and (2) the % of Pos LANs or % of Neg LANs which determines the ratio of two sections in each circle (**D**).

**Second**, were the populations of identified LANs associated with different IVSA events (i.e., ALP, ILP, RNF) separate or overlapping? Our data (**Fig. 8A**) showed a high overlap of LANs between ALP *vs*. RNF in both saline and cocaine mice, but a lower proportion of overlapped LANs between those associated with ALP/RNF *vs*. ILP in cocaine mice, compared to saline mice. Similar to the ratio change between the LAN_Pos_ *vs*. LAN_Neg_, it takes time to develop the separation between ALP/RNF *vs*. ILP of M2 neuronal population in cocaine mice. This separation was observed in cocaine mice, attributed to the reduced neuronal population associated with ILP, during the last 15 min but not the first 15 min on Day 1, and it persisted in both 15 min sessions on Day 5. Thus, we conclude that cocaine IVSA leads to an increase in ALP-but-not-ILP associated neurons and a decrease in ILP-but-not-ALP associated neurons in M2. This suggests a progressive engagement of M2 neurons in cocaine-taking behaviors (i.e., ALP). Furthermore, the separation can be primarily attributed to the frequency (**Fig. 8B**), but not the amplitude (**Fig. 8C**), of the Ca^2+^ transients in M2. We conclude that there is a rapidly developed engagement of M2 neurons in cocaine mice distinguishing ALP *vs*. ILP, which is preserved in the following IVSA days.

**Figure 8.**
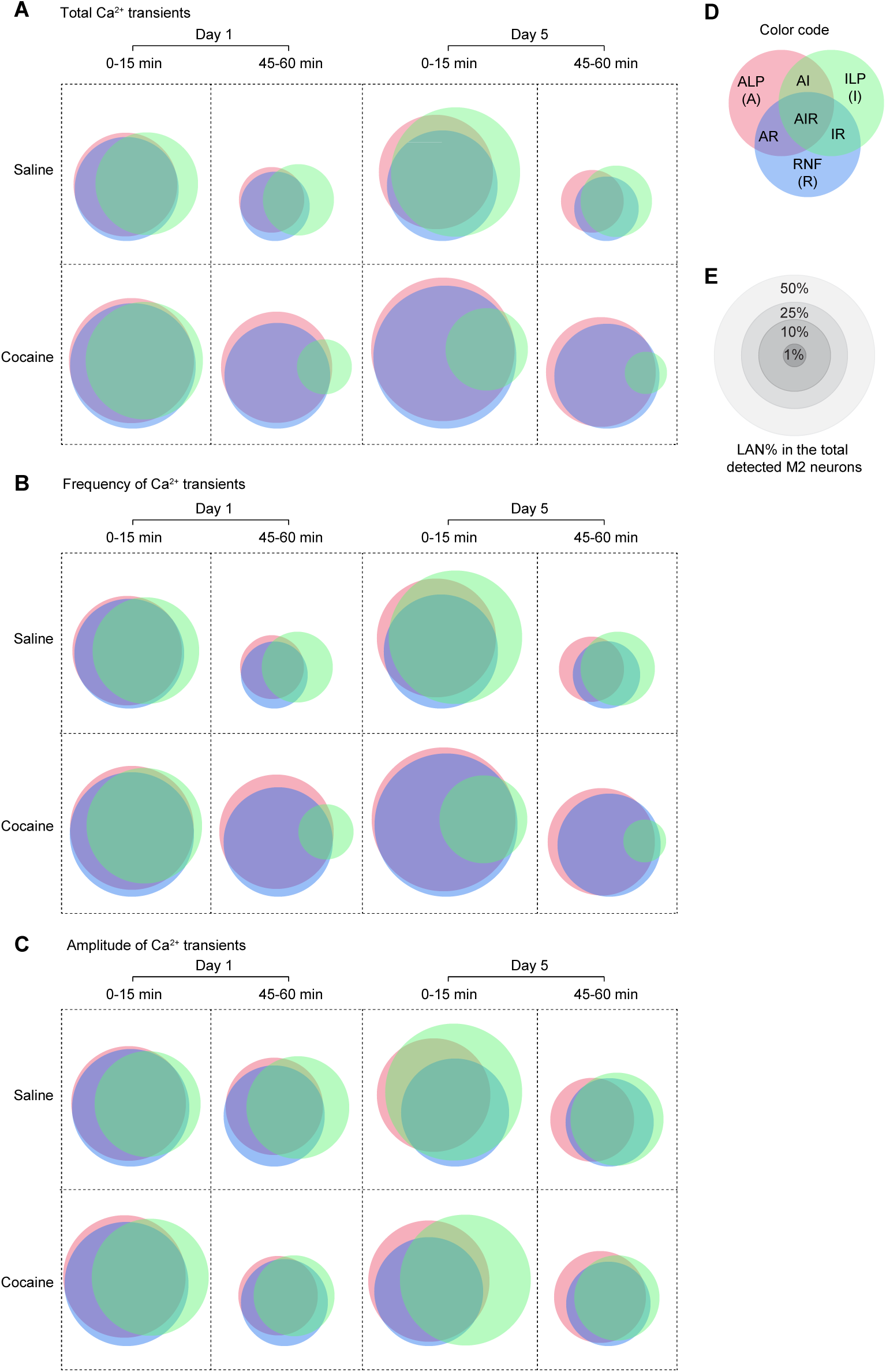
Distribution of IVSA event-associated LANs in M2. **A-C**, Proportional Venn diagrams showing the distribution of M2 neurons with linear association to one or multiple *in vivo* events (i.e., ALP, ILP, or RNF) during the first and the last 15 min of the one-hour saline (upper panels) and cocaine (lower panels) IVSA procedures on Day 1 and Day 5. Ca^2+^ transient quantification was made by the Ca^2+^ transient (i.e., the sum of the amplitude) (**A**), the frequency of the Ca^2+^ transients (**B**), and the amplitude of the Ca^2+^ transients (i.e., the average amplitude) (**C**) in each 30-sec block, which was analyzed by linear regression with the cumulative # of *in vivo* events (i.e., ALP, ILP, RNF) in the corresponding 30-sec block in each 15-min session from individual M2 neurons. **D, E,** Figure legends showing the color codes for different IVSA events and their overlaps (**D**) and the scale of circle size representing the % of LANs (**E**).

## DISCUSSION

This study is the first to identify, in real time, the individual and ensemble neuronal engagement of cortical M2 neurons during drug-taking behaviors. In the present study, we analyzed Ca^2+^ transients by three different readouts: (1) the total Ca^2+^ transients and their (2) frequency and (3) amplitude. The first and last 15-min time windows were used on both Day 1 and Day 5 for behavioral and neuronal data analyses. Special attention was given to the 3-min time window right before ALP/ILP or after RNF. The sequence of IVSA events was also considered. Combing all these data together, a comprehensive understanding of M2 neuronal adaptations during IVSA sessions was further illustrated using the Model events and Model sessions for both saline and cocaine mice in **Fig. 9**. During the first 15 min on Day 1, cocaine mice exhibited a larger Neg neuronal population before ALP and after RNF, suggesting relatively higher Ca^2+^ influx at the time of ALP/RNF, as shown by the model event in **Fig. 9C**. However, during the last 15 min block on Day 1, further consolidated over the entire session on Day 5, cocaine mice show higher Neg neuronal population before ALP but higher Pos neuronal population after RNF, indicating consistently increased Ca²⁺ influx during this 6-minute time window. In addition, the significantly increased LAN_Pos_ neuronal population in each of the 15-min session on the later stage on Day 1 and both early and late stages on Day 5, cocaine mice appear to have a progressively increased Ca^2+^ influx to M2 neurons as indicated by the Model Sessions in **Fig. 9**, which explained the general increase of Ca^2+^ transients in cocaine at the later training stage (**Fig. 2**) based on the data from IVSA -associated 3-min activity before and after ALP/RNF (explained by Model event in **Fig. 9A, C**) and the consecutive IVSA events in the 15 min session (explained by Model session in **Fig. 9B, D**).

**Figure 9.**
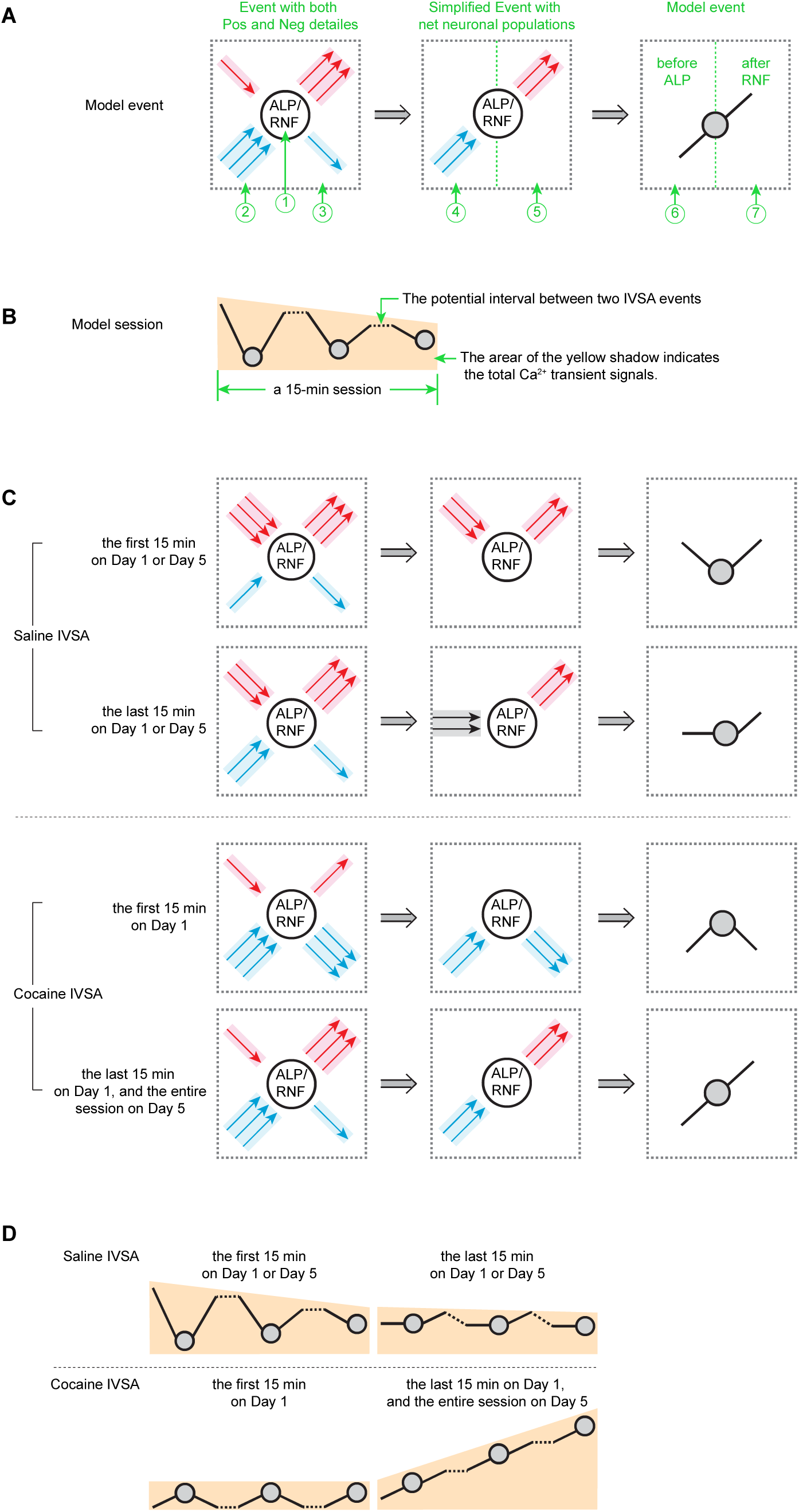
Alterations in Ca^2+^ transients during 15-min sessions of sequential IVSA events. **A**, **How to establish the Model event**: Using the ALP, immediately followed by an IV delivery, as a representative IVSA event, **model event** is derived in three steps. **First**, collecting neuronal activity during an IVSA event. The IVSA event consists of distinct neuronal populations before and after ALP/RNF, as shown in the left panel. The total Ca^2+^ transient-based neuronal categorization is used. Relative to that at the time point of ALP/RNF, Pos neurons, represented by red arrowheads, have higher Ca^2+^ transient signals, and Neg neurons, represented by blue arrowheads, have lower Ca^2+^ transient signals. NR neurons are not considered due to their consistently low level (Fig. 3**-5, S1-S6**). The relative number of Pos and Neg neurons is depicted by the number of arrowheads, with a total of four arrowheads representing a complete set of Pos and Neg neurons. Specifically, one, two, or three arrowheads represent less than 50%, approximately 50%, and greater than 50% of the neuronal population, respectively. **Second**, calculating net neuronal populations. The net neuronal population before and after ALP/RNF is calculated as the balance between Pos and Neg neurons, as shown in the middle panel. **Third**, Model event abstraction. The model event is further abstracted by representing the net neuronal population with a rising and decaying trajectory before and after ALP/RNF, illustrated in the right panel. ①, Time point of ALP, immediately followed by IV delivery as RNF. ②, Pos *vs*. Neg neuronal population before ALP, based on data in Fig. 3I (saline) and **3J** (cocaine). ③, Pos *vs*. Neg neuronal population after RNF, based on data in Fig. 5I (saline) and **5J** (cocaine). ④, Net neuronal population before ALP, balanced from Pos *vs*. Neg neurons at ②. ⑤, Net neuronal population after RNF, balanced from Pos vs. Neg neurons at ③. ⑥, Stroke direction corresponding to the net neuronal population before ALP at ④. ⑦, Stroke direction corresponding to the net neuronal population after RNF at⑤. **B**, **How to establish the Model session**: Model session shows the cumulative changes of Ca^2+^ transient signals across multiple IVSA events (i.e., ALP/RNF) in a 15-min session, consisting of three similar model events (as shown in **A**), and two intervals between IVSA events. The vertical position, or height, of each ALP/RNF spot reflects the the LAN_Pos_ vs. LAP_Neg_ ratio as shown in Fig. 7, with upward shifts occurring when LAN_Pos_% is higher and downward shifts when LAN_Pos_% is lower. The size of the yellow shadow area correlates with the average total Ca^2+^ transients during a 15-min session, as reported in Fig. 2L. **C, Four model events** established for saline IVSA (top two rows) and cocaine IVSA (bottom two rows). **D**, **Four model sessions** established for 15-min sessions of saline IVSA from the first (top row, left) to the last (top row, right) 15 min on both Day 1 and Day 5, and cocaine IVSA from the first 15 min on Day 1 (bottom row, left) to the last 15 min on Day 1 and the entire Day 5 (bottom row, right).

### What do the changes of Ca^2+^ transients mean?

It is widely accepted that the changes in Ca^2+^ transients detected by *in vivo* Ca^2+^ imaging using miniScopes are fundamental processes and *bona fide* indicators of neuronal activation. These transients are different from, but closely related to action potentials (APs). Both the Ca^2+^ transient and the AP are triggered by depolarization of the neuronal membrane, although the source and timing of depolarization differ between the two. Both result in changes in membrane potential. However, they differ in their underlying mechanisms, functions, and propagation characteristics. The neuronal Ca^2+^ transients, as a direct indicator of intracellular signaling and modulation of cellular processes, could be directly triggered by APs, but not all APs are associated with a Ca^2+^ transient as reported before. For example, the influx of Ca^2+^ during each AP could be lower than the threshold of Ca^2+^ influx to be detected as Ca^2+^ transients by miniScope(Oikonomou et al., 2021), thus there is a minimal number of APs within a certain time window to be reached for triggering a Ca^2+^ transient. We call this group of APs corresponding to an individual Ca^2+^ transient as an AP train. It could be quantified by the amplitude of Ca^2+^ transients. Two adjacent but distinct Ca^2+^ transients are assumed to be generated by (1) two trains of APs with a long-enough interval and/or (2) a continuous burst of APs associated with a refractory period during which no further Ca^2+^ influx occurs as a consequence of Ca^2+^ accumulation intracellularly. Thus, the amplitude and frequency of Ca^2+^ transients are related to APs but not paced exclusively by APs, in other words, there is no precise correspondence between APs and Ca^2+^ transients.

When differences in the total Ca^2+^ transients were detected, the frequency and amplitude were also examined to identify the underlying contributors of changes in total Ca^2+^ transients. For example, we found that the acute within-session adaptations of Ca^2+^ transients in saline mice were primarily associated with temporary reductions of Ca^2+^ transient frequency in each daily session, whereas the chronic adaptations in cocaine mice were associated with both increased frequency and amplitude of Ca^2+^ transients. Second, both the frequency and the amplitude of Ca^2+^ transients contributed to the changes of M2 neuronal category based on the total Ca^2+^ transients before ALP and ILP in both saline and cocaine mice. However, the differences between saline *vs*. cocaine mice in their response to RNF is primarily attributable to the frequency but not the amplitude of Ca^2+^ transients. Third, the alterations of LAN% and the separation of LANs between ALP/RNFS vs. ILP in cocaine mice are primarily associated with the frequency of Ca^2+^ transients in M2. Thus, the frequency of the Ca^2+^ transients in M2 neurons is the primary contributor, although not the exclusive one, involved in the drug taking behaviors and also responded to the RNF. Building on our previous discussion of the relationship between Ca^2+^ influx and the APs, we propose that cocaine IVSA modulates the M2 neuron activity by affecting multiple factors, including (1) the number of AP trains, (2) the Ca^2+^ influx associated with each AP train, and (3) the susceptibility of M2 neurons to establish the refractory period during which no further Ca^2+^ influx occurs even if APs continue. We also propose that these effects of cocaine IVSA contribute to the changes in future IVSA behaviors and neuronal responses of M2 to subsequent cocaine IV deliveries.

### M2 neurons are differentially involved in saline vs. cocaine IVSA

Traditional neurobiological studies, which either lacked a longitudinal perspective across days or ignored the changes within a daily session, have often overlooked changes or reported few alterations *in vivo* or *ex vivo* in the saline group under IVSA procedures (Martellotta et al., 1998; Robinson et al., 2022). Interestingly, we found that saline mice had a significant behavioral adaptation within the 1-hr daily sessions on both Day 1 and Day 5, i.e., reduction of the # of ALP, ILP and RNF from the first 15 min to the last 15 min of the 1-hr session on both Day 1 and Day 5. Even more remarkable, M2 seems actively involved in this acute, reversible adaptation of the daily IVSA behaviors in saline mice. We found the within-session alterations of Ca^2+^ transient features across the different IVSA days in saline mice, including (1) the reduction of the general Ca^2+^ transients, particularly the frequency, (2) the decrease of Pos_only_% and increase of Pos_mix_% before ALP, (3) the reduction of the ALP/ILP/RNF associated CML LAN%, (4) the ratio reduction between the ALP/ILP/RNF associated positive *vs*. negative LANs, and (5) the separation of the ALP/RNF associated LANs *vs*. ILP associated LANs. Thus, M2 neuronal activity is significantly reshaped before repeated saline-taking behaviors during the 1-hr daily session. Our data indicate that early IVSA behaviors may influence M2 neuronal activity later in the session, but this effect is confined within the session and is reversible in subsequent days.

Similar to saline mice, we found within-but not across-day adaptations of the IVSA behaviors in cocaine mice (e.g., the linear increase of IAI in the first 15 min and then remained at the high level in the last 15 min on both Day 1 and Day 5). However, the majority of Ca^2+^ transient-associated adaptations of M2 in cocaine mice are not the same between Day 1 vs. Day 5. Specifically, we observed that some of the altered M2 neuronal features, including the increased general (i.e., IVSA-event non-associated) Ca^2+^ transients, attributable to both higher frequency and larger amplitude), increased Pos_mix_% and decreased Neg_only_% after RNF, reduced CUM LAN% associated with ILP, reduced ratio between positive *vs*. negative LANs associated with ALP and RNF, and separation of LANs associated with ALP/RNF *vs*. ILP, at the late stage on Day 1 were well maintained throughout the 1-hr daily session on Day 5 in cocaine mice. We conclude that, different from the reversible alterations associated with saline IVSA, repeated cocaine IVSA resulted in chronic neuronal adaptations in M2. The similar patterns of cocaine IVSA behaviors on Day 1 and Day 5 are associated with distinct neuronal activity in M2. It is worth noting other adaptation patterns of Ca^2+^ transient features, such as the elimination of within-session fluctuations of % of Pos_only_ and Pos_mix_ and across-day fluctuations of % of Neg_only_ and Neg_mix_ before ALP observed in saline mice, as well as reduced ratio of Neg_only_% *vs*. Neg_mix_% from Day 1 to Day 5. Cocaine IVSA appears to prevent the acute (e.g., within-session) adaptations while promoting the prolonged (e.g., across-days) adaptations.

### M2 neurons are differentially involved in the initiation of ALP *vs*. ILP

From the early to the late stage of IVSA daily sessions on both Day 1 and Day 5, the proportion of three different neuronal clusters based on the relative changes of Ca^2+^ transients (i.e., Pos, NR, and Neg) were consistently maintained before ALP or ILP in both saline and cocaine mice. Lower Pos% and higher Neg% were detected before ALP in cocaine mice, relative to those in saline mice. There were no significant differences of Pos% or Neg% between saline vs. cocaine mice detected before ILP. Thus, the changes observed before ALP cannot be simply attributed to the lever press itself but more related to some unique features associated with ALP. For example, ALP, but not ILP, was associated with cue light presentation and IV delivery. When we looked into the subtypes of neuronal clusters in M2, we identified that consistently higher Neg% before ALP in cocaine mice on both Day 1 and Day 5 should be attributed to the higher Neg_only_% on Day 1 and higher Neg_mixed_% on Day 5, respectively. The changes of subtype neuronal proportions from Day 1 to Day 5 was absent before ILP in cocaine mice, although they established the within-session fluctuation of % of Pos_only_ and Pos_mix_ before ILP which were flat in saline mice. Relative to the neurons categorized as Neg_mixed_, the Neg_only_ neurons are assumed to be better correlated with a consistent reduction of Ca^2+^ influx. Thus, we conclude that synchronized M2 activities seem necessary to initiate the cocaine taking behaviors on Day 1, and a relative lower level of synchronized reduction of Ca^2+^ influx to M2 neurons seems sufficient to initiate the cocaine taking behaviors on Day 5. Our data indicate that the adaptation of M2 neurons by the drug taking experience on early days facilitates the maintenance of persistent cocaine taking behaviors on later days. This real time adaptations observed in M2 may provide a neuronal substrate to explain the transition of drug use to drug mis-use as observed in the laboratory animal model and in clinical human cases (Kuhn et al., 2019; Lees et al., 2021). Furthermore, we found that saline mice had a higher LAN% associated to both ALP and ILP during the first 15 min, compared to the last 15 min on both Day 1 and Day 5. In contrast, cocaine mice showed significantly lower LAN% associated with ILP after 15 min on Day 1, which remained at low levels on Day 5. In cocaine mice, more M2 neurons displayed progressively increased Ca^2+^ transients as they engaged in more ALP behavior. Notably, most of these neurons were exclusively involved in ALP. The initiation of ILP might be related to a distinct and smaller neuronal population within M2.

### M2 neurons differentially responded to IV delivery of saline vs. cocaine

Different from the consistent proportions of all the four subpopulations of M2 neurons after RNF, cocaine mice showed lower Pos_mix_% and higher Neg_only_% during the first 15 min on Day 1, which were adjusted to get close to the levels as in saline mice during the last 15 min on Day 1, and then fully returned to the level as in saline mice on Day 5. The temporary but not persistent inhibition of a relatively larger population of M2 and activation of a smaller neuronal population right after cocaine IV injection could be explained by the re-establishment of homeostasis by compensating the neuronal effects of IV cocaine in M2 after the very early exposure to IV cocaine. Thus, a potential rebound of M2 neuronal responses could occur during the drug withdrawal period. Another potential explanation of time-limited responses to IV cocaine is that M2 neurons were desensitized to IV cocaine after repeated exposure. This may support no further adaptations of M2 activity at the withdrawal stage, which is supported by our recent publication showing no excitability changes in M2 neurons 24 hr after the last cocaine IVSA session (Huang and Ma, 2023a). One more alternative explanation could be a temporary response to the 20-sec cue light presented right after ALP. The potential effects of cue light itself could be partially ruled out, as similar cue light exposure in saline mice did not produce the same neuronal alterations as seen in cocaine mice. The response to IV cocaine might be attributable to the light serving as a conditioned cue signaling drug availability. Therefore, the early alterations in the M2 neural population could reflect a prediction of the reinforcer, which may gradually evolve into a prediction error, a concept established in dopamine neurons as a reward expectation theory(Lerner et al., 2021; Schultz et al., 1997). If this is the case, we would expect no more changes of M2 neuronal population responses after RNF, as the prediction aligned with the expected outcome (i.e., successful IV delivery of cocaine at the expected time window). We would further anticipate an evoked M2 neuronal response following ALP during cocaine seeking test when cocaine is not available as expected.

### Other considerations and insights

#### First, why were IVSA event-associated 30 sec bins used in Ca^2+^ transient analyses?

Although Ca^2+^ imaging data were collected at a high sampling rate (30 Hz), smaller time windows did not necessarily yield greater sensitivity. This is likely due to the slow kinetics of Ca^2+^ transients (with a decay of 5-15 seconds) and their low frequency (0.02-0.07 Hz, or generally no more than two Ca^2+^ transients per 30 seconds. We applied multiple machine learning methods (e.g., Convolutional Neural Network, Recurrent Neural Network, Extreme Gradient Boosting, and Support Vector Machine) to train the models to distinguish the Ca^2+^ transient data across several experimental factors, such as saline *vs*. cocaine, IVSA Day 1 *vs*. Day 5, the first *vs*. the last 15 min of the daily 1-hr session. Our data demonstrated that a bin size of at least 15 seconds was required to detect differences in most experimental scenarios. We also developed a python-based pipeline using evolutionary algorithms to optimize the bin sizes over 100 generations. This approach revealed that a bin size of ∼30 seconds, with six bins per time window (3 minutes total), was among the most sensitive for detecting differences between groups (e.g., saline *vs*. cocaine) and within groups (e.g., Day 1 *vs*. Day 5). It is worth noticing that during the 3-min time window, the neuronal alterations in M2 may not be exclusively attributed to drug taking behaviors. For example, the general locomotion could be another significant factor.

#### Second, would similar responses in M2 neurons be observed in sucrose self-administration model?

Our previous whole-cell patch clamp recordings demonstrated that, different from cocaine IVSA withdrawal, sucrose self-administration withdrawal did not change M2 excitability, which is similar to what was seen in the saline group. Additionally, inhibiting M2 excitability had no effect on sucrose-seeking behavior (Huang and Ma, 2023a). We anticipated that Ca^2+^ transient signals in M2 neurons during sucrose-taking behaviors share features with both the saline and cocaine groups, yet exhibit a unique pattern. This pattern likely reflects a transitional state, characteristic of a natural reinforcer, situated between the saline (non-reinforcer) and cocaine (substance with misuse potential) groups. Specifically, given the reinforcing effects of sucrose pellets, we hypothesize that Ca^2+^ transients in M2 neurons on Day 1 of sucrose-taking will resemble those in cocaine-treated mice on their Day 1. However, due to sucrose’s limited misuse potential, relative to that of cocaine, we expect that within-session changes in Ca^2+^ transients in sucrose mice on Day 1 are reversible, as seen in saline mice. For instance, we predict an increase in Ca^2+^ transient signals from the first 15 minutes to the last 15 minutes during the 1-hour session on both Day 1 and Day 5 in sucrose-treated mice. In conclusion, M2 neurons appear to exhibit temporarily enhanced but not prolonged functional changes associated with sucrose-taking behavior.

#### Third, IT or PT neurons in M2 were detected?

Our *in vivo* Ca^2+^ recordings were conducted primarily in the superficial layers in M2, which mainly contain intratelencephalic (IT), rather than the pyramidal tract (PT) neurons (Hooks et al., 2013; Peters et al., 2017). Unlike PT neurons, which project to brain stem and the spinal cord to directly regulate movement, IT neurons primarily project to other cortical areas and the striatum, responsible for integrating information within the cerebrum. This supports the idea that the detected M2 neurons in addiction behaviors is linked to more complex, integrative functions as mentioned above.

## CONCLUSIONS

M2 neurons are sensitive to and significantly affected by the previous operant behaviors and the history of drug exposure, which in turn sculpt the upcoming operant behaviors and the response to drugs. As critical nodes of the reward and decision-making loop, M2 neurons appear to be the governing center in facilitating the establishment of addiction-like behaviors associated with chronic functional changes.

## MATERIALS AND METHODS

### Experimental subjects, surgical procedures, behavioral test and in vivo Ca2+ imaging recordings

All procedures were performed in accordance with the United States Public Health Service Guide for Care and Use of Laboratory Animals and were approved by the Institutional Animal Care and Use Committee at Indiana University School of Medicine. Eighteen 5-week-old male C57BL/6 were purchased from the Jackson Laboratory and group housed for 5-7 days until surgery and behavioral procedures started (**Fig. 1A**). During these procedures, mice were singly housed on a 12-hour light/dark cycle with free access to chow and water in home cage.

### Surgical procedures

#### (1) Microinjection of AAV

Mice were anesthetized with 2.5% isoflurane for induction and maintained with ∼1.2%. A 28-gauge injection needle was used to unilaterally inject the tAAV1-Syn-jGCaMP8f-WPRE or AAV1-CaMKIIa-jGCaMP8f-WPRE solution (0.5 µl/site, 0.1 µl/min) *via* a Hamilton syringe into the M2 (coordinates in mm: AP, +1.80; ML, ±0.60; DV, −1.30 for M2), using a Pump 11 Elite Syringe Pumps (Harvard Apparatus). Injection needles were left in place for 5 min following injection.

#### (2) Lens implantation

A couple of minutes after withdrawing the AAV injection needle, a unilateral GRIN lens (Inscopix Inc, #1050-004595, Diameter: 1.0 mm; length: ∼4.0 mm; Working Distance: 200 µm) was lowered through the cranial window to 200 µm above the center of the virus injection site. The open space between the lens’ side and the skull opening was sealed with surgical silicone (Kwik-Sil) and secured by dental cement (C&B Metabond). The exposed part of the lens above the skull was further coated with black cement (Lang Dental Mfg. Co.’ Inc.).

#### (3) Catheter implantation

3∼4 weeks later, catheter implantation was made as described previously(Guo et al., 2022; Huang and Ma, 2023a; Ma et al., 2012; Ma et al., 2014; Ma et al., 2016). Briefly, a silastic catheter was inserted into the right jugular vein, and the distal end was led subcutaneously to the back between the scapulae. Catheters were constructed from silastic tubing (length, 55 mm; inner diameter, 0.3 mm; outer diameter, 0.6 mm) connected to a One Channel Vascular Access Button^TM^ for Mice (Instech Labs). Aluminum VABM1C cap (Instech Labs) was attached to protect the button. The catheter was flushed daily after implantation with 1 ml/kg body weight of heparin (10 U/ml) and gentamicin antibiotic (5 mg/ml) in sterile saline to help protect against infection and catheter occlusion.

#### (4) Base-plating

After the catheter implantation, the mouse was maintained anesthetized with isoflurane. The cement on top of the GRIN lens was carefully removed using drill bits until the lens was exposed. The top of the lens was then cleaned using lens paper and a cleaning solution. A metal baseplate was mounted onto the skull over the lens using Loctite super glue gel, guided by a MiniScope for optimal field of view. Once the baseplate was securely mounted, the MiniScope was removed. A protective cap was attached to the baseplate, and the mouse was returned to its home cage.

#### (5) Verification of AAV expression and the lens location

After the completion of the behavioral tests, M2-containing coronal slices were prepared as described before(Huang and Ma, 2023b), then fixed in 4% Paraformaldehyde (PFA) for no less than a couple of hours. After a brief rinse with PBS, slices were mounted with Prolong^TM^ Gold antifade mounting reagent with DAPI (Invitrogen, Cat# P36931). Confocal imaging was performed using a Zeiss LSM 800 confocal microscope. The animals included in this study were all approved with (a) highly enriched AAV expression in mostly pyramidal neurons, within-M2 viral injection site, and (b) the footprint of the GRIN lens tip at the top of M2 (**Fig. 2A**).

### Open field test

∼6 days after the catheter implantation, mice were placed in a Black Plexiglas testing chamber (Maze Engineers; 60 cm for both side-widths x 50 cm in height with a floor illumination at ∼60 lux) (Shan et al., 2019) (**Fig. 1A**). Mice were placed in the peripheral area facing one of the randomly assigned sidewalls and then allowed to freely explore the open field. Total testing time was 5 min. Behavioral videos were processed by EthoVision XT14 Software (Noldus) and the total distance traveled during the 5 min period was calculated.

### Intravenous self-administration

#### (1) Self-administration apparatus

Experiments were conducted in operant-conditioning chambers enclosed within sound-attenuating cabinets (Med Associates). We randomly assigned 8 mice to the saline control group and 10 mice to the cocaine treatment group. Each chamber contained two levers, randomly assigned as either active or inactive, a food dispenser, and the conditioned stimulus light (i.e., cue light) 2 cm above each lever. No lab chow or water was provided in the chambers during the training or testing sessions.

#### (2) Intravenous cocaine self-administration training

∼7 days after catheter implantation, cocaine self-administration training began with an overnight session at 7 PM to 7 AM the following day. The daily training session, consisting of one hour per day for five days, commenced the day after the overnight session. The same training protocol was used in overnight and daily sessions. Mice were placed in the self-administration chamber on a fixed ratio (FR) 1 reinforcement schedule with the house light on. Active lever presses (ALP) resulted in a cocaine infusion (0.75 mg/kg over ∼2 sec) and illumination of a cue light above the active lever for 20 sec with the house light off. In contrast, the inactive lever presses (ILP) led to no outcome (i.e., neither IV infusion nor cue light presentation) but were also recorded. Mice that received at least 60 cocaine rewards in the overnight session were allowed to move to daily self-administration of cocaine ∼24 hr after the overnight training on an FR1 reinforcement schedule. All mice (n=18) transitioned to daily training sessions after just one overnight training session.

### *In vivo* Ca^2+^ imaging recording during cocaine self-administration

Mice were habituated to the *in vivo* Ca^2+^ recording procedure by mounting the miniScope V4 (OpenEphys) to the pre-anchored base plate and recording for 5 min per day at home cage for 3 days before starting the IVSA procedures. Our data presented in **Figs. 2-8, S1-S7** were collected during the 1-hr IVSA daily sessions on the first and the last IVSA training days (*i.e.*, Day 1 and Day 5). Facilitated by the customized commutator from Doric, mice were allowed to self-administer cocaine *via* pre-implanted IV catheter with no interruptions from double lines (i.e., the IV solution line and the miniScope coax cable). Data Acquisition (DAQ) box, supported by an open source, C^++^ and Open Computer Vision (OpenCV) libraries-based software, were used to collect both Ca^2+^ and behavioral video streams simultaneously controlled by the operant chamber software, MED-PC (Med Associates) *via* a TTL adaptor. The sampling frequency was 30 Hz.

### Ca^2+^ imaging analyses

#### (1) Extraction of Ca^2+^ transient traces from the raw videos

Among multiple types of computational tools established previously to extract Ca^2+^ transients from raw videos, a Python-based analysis pipeline, Minian(Dong et al., 2021), was used in our data analyses due to its low memory demand and user friendly parameter options. In brief, there were five steps in the pipeline. First, multiple raw videos were batch loaded and subjected to a *PREPROCESSING* stage, where sensor noise and background fluorescence from scattered light were removed. Second, rigid brain motions were corrected by *MOTION CORRECTION*. Third, the initial spatial and temporal matrices for later steps were generated by a seed-based approach, called *SEEDS INITIALIZATION*. Fourth, the spatial footprints of cells were further refined. Fifth, the temporal signals of cells were also refined. The last two steps, the *SPATIAL UPDATE*, and the *TEMPORAL UPDATE*, as the core computational components based on CNMF algorithm, were repeated at least one more time.

#### (2) Cell verification

Our team has developed an open-source GUI using the python package PYQT5 for cell verification and Ca^2+^ transient identification. Synchronized visualization *via* a user-friendly single window allowed us to evaluate the CNMF output comprehensively. We visualized multidimensional Ca^2+^ signals generated by CNMF variables “C” or “S”, and directly calculated by our GUI tool such as raw fluorescent signals (F), filtered fluorescent signals (F_filtered_), noise, signal-to-noise ratio (SNR), and the relative change of fluorescent intensity (ΔF/F). The individual cell footprint was generated by the CNMF variable “A” on top of the raw Ca^2+^ video or the CNMF-processing Ca^2+^ video. Selection of the footprint or the cell ID from the cell ID listed achieved by CNMF was directly associated with the raw image, the processed image, the footprint, the cell ID and the Ca^2+^ transient traces. Cells were excluded if any of this information appeared artifactual. We also plotted the SNR values for each cell. Cells with low SNR values (e.g., SNR<2) were excluded unless the Ca^2+^ transients were identifiable. There were 2-20 % of CNMF-identified cells per recording session per animal excluded from further data analyses.

#### (3) Ca^2+^ transient identification

**First**, we used the Ca^2+^ transient kinetics-based auto-identification module in our newly developed GUI platform. The transient was detected by the CNMF variable “C” and the ΔF/F. <u>Ca^2+^ transient amplitude</u> was determined by the peak value of the ΔF/F. The minimal Ca^2+^ transient amplitude was set as ΔF/F = 1, which required that the increase of the fluorescent intensity is at least 100% from the background level. The minimal inter-transient interval was set at 45 frames or >1.5 sec. **Second**, we checked the Ca^2+^ transient traces and further adjustment was manually made to accept new Ca^2+^ transients or reject the auto-identified Ca^2+^ transients. **Lastly**, we established machine-learning modules by loading the data sets of Ca^2+^ traces with auto/manually identified Ca^2+^ transients for the training, validation, and testing. Briefly, two variants of Recurrent Neural Networks machine learning models (i.e., LSTM and GRU) and two versions of the Transformer model (i.e., the original model and one using local attention) were tested. Our data demonstrated that GRU-based machine learning model is highly reliable, achieving an F1 score (i.e., a statistical measure of predictive performance) of over 0.9 (e.g., F1 = 0.94). This performance was attained using a limited dataset consisting of Ca^2+^ traces from only 10-20 cells over a 15-min window to train the machine learning model.

#### (4) Bin-sized Z score during a 15-min session

A bin of 30 seconds was used in preparing data for **Figs. 3-8, S1-7**. For those spanning across two adjacent 30-sec blocks, the Ca^2+^ transients were included in the 30-sec block where they started. Three readouts of the Ca^2+^ transient signals were used for their quantification in each 30-sec block:

<u>Total Ca^2+^ transients per bin</u> were calculated by the summation of the amplitude of all the Ca^2+^ transients in each 30-sec block of the observed neurons.

<u>Ca^2+^ transient frequency per bin</u> was calculated by the number of Ca^2+^ transients in each 30-sec block of the observed neurons divided by 30 sec. The unit is Hz.

<u>Ca^2+^ transient amplitude per bin</u> was calculated by the average of the amplitude of the Ca^2+^ transients in each 30-sec block of the observed neurons.

Z score in each 30-sec block during a 15-min time window for each neuron was calculated by the following formula:

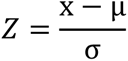

In which x is a specific Ca^2+^ transient readout (i.e., total Ca^2+^ transient, Ca^2+^ transient frequency, or Ca^2+^ transient amplitude) from an observed neuron in any of the thirty 30-sec time blocks, µ and σ are the corresponding mean value and the standard deviation of the thirty 30-sec blocks of the observed neuron.

#### (5) Bin-sized ΔZ score within 3 min before ALP/ILP or after RNF during a 15-min time window

##### Neuronal ΔZ score of the total Ca^2+^ transients in a 30-sec block

Using the ALP-associated ΔZ scores of the total Ca^2+^ transients in **Fig. 3 C, D** as examples, each ΔZ score of the total Ca^2+^ transient in a 30-sec block was calculated by the total Ca^2+^ transient Z score minus the Z score in the bin where ALP occurred. Neurons from an animal which had at least 1 ALP during a 15-min time window were included in calculating the ALP-associated ΔZ. Assuming there were n (as the number of) ALPs during the 15-min time window (ALP#1, ALP#2, …., ALP#n), then we would have n sets of six ΔZ values for each of the six 30-sec block with 3 min before ALP. The averaged six ΔZ values within 3 min before ALP from the n sets are presented in **Fig. 3 C, D**, in which each row of the six blocks represented the neuronal changes of the total Ca^2+^ transients in each 30-sec block within 3 min before ALP. Similarly, the neuronal changes of the total Ca^2+^ transients in each 30-sec block within 3 min before ILP are shown in **Fig. 4 C, D**, and the neuronal changes of the total Ca^2+^ transients in each 30-sec block within 3 min after RNF are presented in **Fig. 5 C, D**.

##### Neuronal ΔZ score of the frequency of Ca^2+^ transients in a 30-sec block

Similar calculations as the neuronal ΔZ score of the total Ca^2+^ transients described above, the Z scores of the frequency of the Ca^2+^ transients in each of the six 30-sec blocks was used in ΔZ score quantification. Specifically, the neuronal changes of the frequency of Ca^2+^ transients in each 30-sec block within 3 min before ALP are shown in **Fig. S1 A**, **B,** the neuronal changes of the frequency of Ca^2+^ transients in each 30-sec block within 3 min before ILP are presented in **Fig. S3 A**, **B**, and the neuronal changes of the frequency of Ca^2+^ transients in each 30-sec block within 3 min after RNF are shown in **Fig. S5 A**, **B**.

##### Neuronal ΔZ score of the amplitude of Ca^2+^ transients in a 30-sec block

Similar calculations as the neuronal ΔZ score of the total Ca^2+^ transients described above, the Z scores of the average amplitude of the Ca^2+^ transients in each of the six 30-sec blocks was used in ΔZ score quantification. Specifically, the neuronal changes of the amplitude of Ca^2+^ transients in each 30-sec block within 3 min before ALP are illustrated in **Fig. S2 A**, **B,** the neuronal changes of the amplitude of Ca^2+^ transients in each 30-sec block within 3 min before ILP are shown in **Fig. S4 A**, **B**, and the neuronal changes of the amplitude of Ca^2+^ transients in each 30-sec block within 3 min after RNF are shown in **Fig. S6 A**, **B**.

#### (6) ΔZ score-based sorting of the 30-sec bin

Each of the IVSA events (i.e., ALP, ILP, RNF)-associated 30-sec blocks before ALP/ILP or after RNF are categorized as positive response if ΔZ is more than +0.1, no response if ΔZ is between −0.1 to +0.1, or negative response (if ΔZ is less than −0.1). Specifically, the ΔZ score-based sorting of the 30-sec bins are provided in panels **E-H** in **Figs. 3-5**, and panels **E-F** in **Figs. S1-S6**.

#### (7) ΔZ score-based neuronal sorting of the 3-min time window

The IVSA event (i.e., ALP, ILP, or RNF)-associated 3-min time window for each neuron consisted of six 30-sec blocks. The 3-min window associated with ALP or ILP, which means the 6^th^ 30-sec block (i.e., −30 to 0 sec or −0.5 to 0 min) was the block when the ALP or ILP occurred, was categorized as (a) positive (Pos), (b) non-responsive (NR), and (c) negative (Neg) if the 5^th^ 30-sec block (i.e., −60 to −30 sec or −1 to 0.5 min, representing the 30-60 sec before ALP) was classified as positive, non-responsive, and negative, respectively. The 3-min window associated with RNF, which means the 1^st^ 30-sec block (i.e., 0 to 30 sec or 0 to 0.5 min) was the block when the RNF occurred, was categorized as positive (Pos) and negative (Neg) if the nearest responsive 30-sec block (i.e., a bin defined as either positive or negative response as described in “ΔZ score-based sorting of the 30-sec bin”) was classified as positive and negative, respectively. The 3-min window associated with RNF was categorized as non-responsive (NR) if all the five 30-sec blocks during 30 to 180 sec (i.e., 0.5 to 3 min) are sorted as no responsive.

% of animal event (i.e., ALP, ILP, or RNF)-associated neuronal population in each 15-min was calculated for each experimental mouse as below.

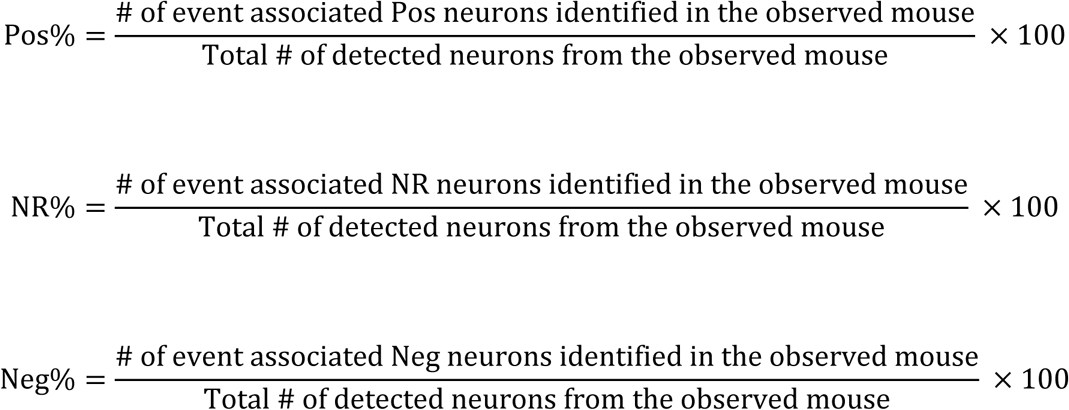

Thus, Pos%+ NR%+ Neg% = 100% for each mouse. The % of ALP-associated neuronal population in saline *vs*. cocaine mice is shown in **Fig. 3I-M**. The % of ILP-associated neuronal population in saline *vs*. cocaine mice is shown in **Fig. 4I-M**. The % of RNF-associated neuronal population in saline *vs*. cocaine mice is shown in **Fig. 5I-M**.

#### (8) Subcategories of Pos and Neg neuronal populations

The Pos neurons were further categorized into Pos_only_ (*i.e*., all the five 30-sec blocks before ALP/ILP or after RNF were identified as Pos) *vs*. Pos_mixed_ (*i.e.*, at least one of the five 30-sec blocks before ALP/ILP or after RNF were identified as NR or Neg). Similarly, Neg neurons were also further categorized into Neg_only_ (*i.e*., all the five 30-sec blocks before ALP/ILP or after RNF were identified as Neg) *vs*. Neg_mixed_ (*i.e.*, at least one of the five 30-sec blocks before ALP/ILP or after RNF were identified as NR or Pos). The % of detected M2 neurons in each subcategory was calculated for each animal similarly. For example, ALP-associated Pos_mixed_ % for a mouse was calculated by the # of ALP-associated Pos_mixed_ neurons identified in the observed mouse divided by the total # of detected M2 neurons from the observed mouse. The % of ALP-associated neuronal sub-population (i.e., only vs. mixed) in saline *vs*. cocaine mice is shown in **Fig. 3N-Q**. The % of ILP-associated neuronal sub-population (i.e., only vs. mixed) in saline vs. cocaine mice is shown in **Fig. 4 N-Q**. The % of RNF-associated neuronal sub-population (i.e., only vs. mixed) in saline vs. cocaine mice is shown in **Fig. 5 N-Q**.

#### (9) Linear regression analysis

The linear regression analysis was performed between (1) the Z scores of the total Ca^2+^ transients (**Figs. 6, 7, 8A,B, S7**), the frequency of Ca^2+^ transients (**Fig. 8C**), or the amplitude of Ca^2+^ transients (**Fig. 8C**) and (2) the # of ALP, ILP or the RNF in each of the fifteen 30-sec blocks during a 15-min session. A linear IVSA-event (i.e., ALP, ILP, or RNF)-associated neuron (denoted as LAN) was identified when the p value of the F test for the linear regression was less than 0.05, indicated the regression is non-zero, and the p value of the Pearson correlation analysis was also less than 0.05. The # of ALP, ILP and RNF was set up in two different ways, including (1) the counts of ALP, ILP or RNF during each 30-sec block, and (2) the counts of ALP, ILP or RNF in not only the current 30-sec block but also the previous 30-sec block(s) if there was any within the 3-min time window. Thus, we performed and compared two types of linear regression analyses, named non-cumulative (nonCML) linear regression, and cumulative (CML) linear regression, respectively as shown in **Fig. 6**. Only CML linear regression-identified LANs were further analyzed in **Figs. 7, 8, S7**.

### Statistical Analysis

Data were collected from 8 saline mice and 10 cocaine mice *in vivo,* shown as mean ± SEM (except in **Fig. 2I-K** where three dashed lines split each violin plot at the quartile positions) and/or individual values from each inter-ALP-interval (**Fig. 1R-U**), each neuron (**Figs. 2C-H**), or each mouse (**Figs. 1C-K, 2K-N, 3I-Q, 4I-Q, 5I-Q, 6A-L**). Using GraphPad Prism 10, statistical significance was assessed by Student’s t test (**Fig. 1B**), two-way ANOVA (**Figs. 1C-N; 3-5 I-Q; 6 A-L**), three-way ANOVA, followed by Bonferroni post-hoc tests (**Fig. 2I-N**), or simple linear aggression and Pearson correlation analysis (**Fig. 1R-U** and neuronal characterization in **Figs. 6-8**). Statistical significance was considered to be achieved if p < 0.05.

## ACKNOWLEDGEMENTS

This work was supported by NIH grants (R01AG072897, R01AA025784, and R01DA059548). We thank the support from High-Performance Computing and Storage at Indiana University.

## AUTHOR CONTRIBUTION

Experimental design: YC, CG, YYM. Data collection: YC. Data analysis: YC, HF, AK, MAL, CR, DL, SF, CG, YYM. Result discussion: YC, HF, AK, MAL, CR, LT, DL, SF, CG, YYM. Manuscript preparation: YC, HF, AK, MAL, CR, LT, DL, SF, CG, YYM.

