## SUPPLEMENTARY FIGURES 1-7 and Tables for "Decoding Secondary Motor Cortex Neuronal Activity during Cocaine Self-Administration: Insights from Longitudinal *in vivo* Calcium Imaging"

#### SUPPLEMENTARY FIGURES & LEGENDS

#### Figure S1

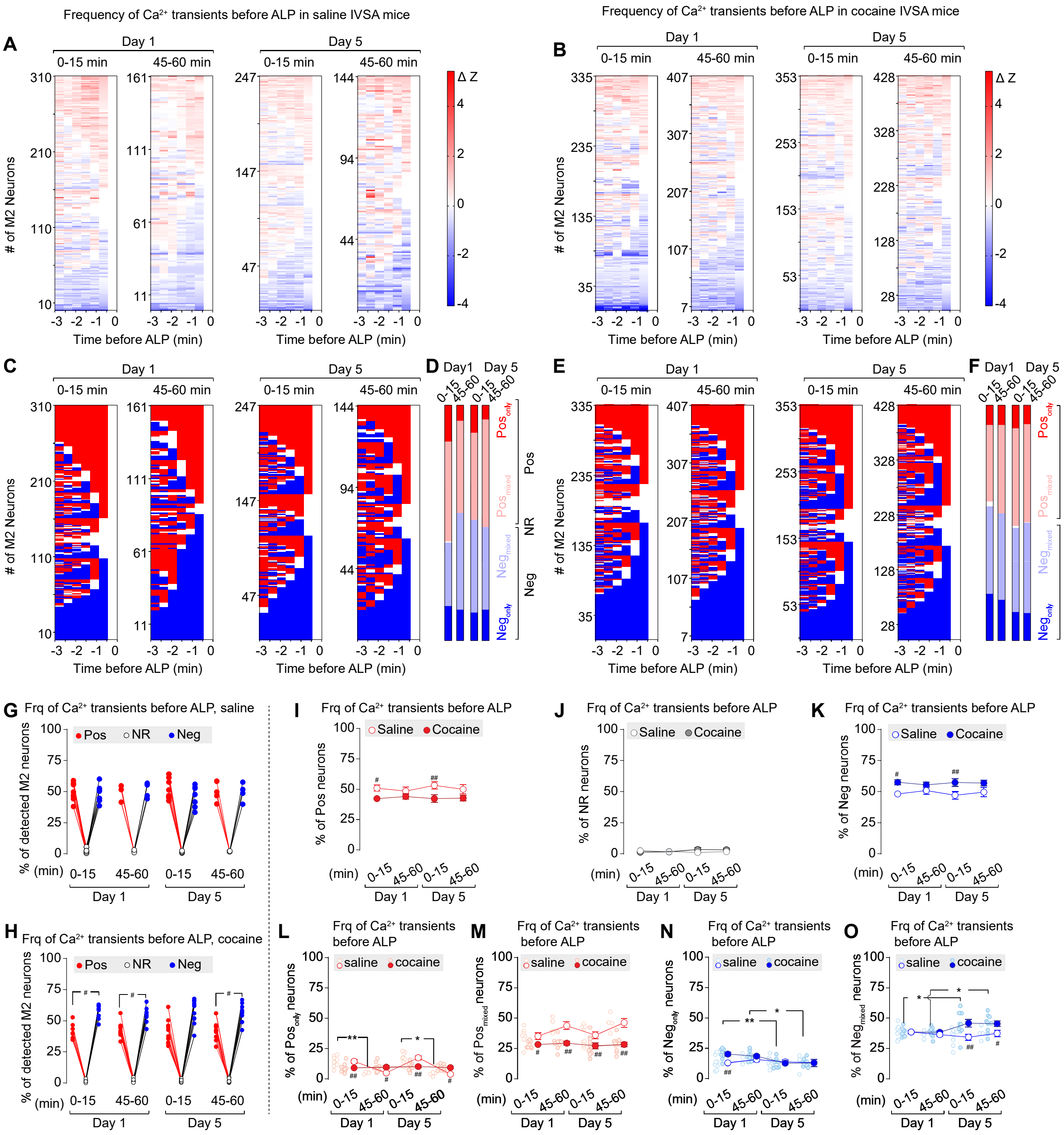

Frequency of Ca2+ transients before ALP in saline IVSA mice
Frequency of Ca2+ transients before ALP in cocaine IVSA mice
A
B
C
D
E
F
Day 1
Day 5
0-15 min
45-60 min
### of M2 Neurons
Time before ALP (min)
Δ Z
Pos only
Pos mixed
Neg mixed
Neg only
Pos
NR
Neg
G
Frq of Ca2+ transients before ALP, saline
H
Frq of Ca2+ transients before ALP, cocaine
% of detected M2 neurons
I
J
K
Frq of Ca2+ transients before ALP
Saline
Cocaine
% of Pos neurons
% of NR neurons
% of Neg neurons
L
M
N
O
Frq of Ca2+ transients before ALP
saline
cocaine
% of Pos only neurons
% of Pos mixed neurons
% of Neg only neurons
% of Neg mixed neurons
(min)

**Figure S1. Frequency of $Ca^{2+}$ transients in M2 neurons during 3 min before ALP in saline vs. cocaine mice.**

**A**, **B**, Heat maps of ΔZ scores of $Ca^{2+}$ transient frequency in detected M2 neurons displayed by rows of six 30-sec blocks within 3 min before ALP in the first (left in each pair of the trace columns) and the last (right in each pair of the trace columns) 15 min during Day 1 and Day 5 IVSA sessions in saline (**A**) and cocaine (**B**) mice.

**C-F**, $Ca^{2+}$ transient frequency ΔZ based-classification of each 30-sec block of M2 neurons within 3 min before ALP in the first (left in each pair of the trace columns) *vs*. the last (right in each pair of the trace columns) 15 min during Day 1 and Day 5 IVSA sessions in saline (**C**) and cocaine (**D**) mice. The proportions of M2 neurons in each category in a 15-min session are summarized in **E** for saline mice and **F** for cocaine mice.

**G**, **H**, Summarized data showing the percentage of three ALP & $Ca^{2+}$ transient frequency-based categories of M2 neurons in saline (**G**) and cocaine (**H**) mice. Both saline (**G**, neuron type × IVSA training session interaction $F_{6,40}=0.3$, $p=0.91$; neuron type $F_{1,20}=279.5$, $p<0.01$; and IVSA training session $F_{3,20}=52.8$, $p<0.01$) and cocaine (**H**, neuron type × IVSA training session interaction $F_{6,66}=0.1$, $p=0.99$; neuron type $F_{1,35}=513.3$, $p<0.01$; and IVSA training session $F_{3,33}=308.9$, $p<0.01$) mice demonstrated different proportions among three neuronal types.

**I-K**, Summarized data showing smaller proportions of ALP & $Ca^{2+}$ transient frequency-categorized Pos neurons (**I**, saline/cocaine × IVSA training session interaction $F_{3,53}=0.4$, $p=0.75$; saline/cocaine $F_{1,53}=13.7$, $p<0.01$; and IVSA training session

$F_{3,53}$=0.1, p=0.94), similar proportions of ALP-categorized NR neurons (**J**, saline/cocaine × IVSA training session interaction $F_{3,53}$=0.9, p=0.47; saline/cocaine $F_{1,53}$=0.5, p=0.50; and IVSA training session $F_{3,53}$=0.1, p=0.95), and larger proportions of ALP-categorized Neg neurons (**K**, saline/cocaine × IVSA training session interaction $F_{3,53}$=0.4, p=0.78; saline/cocaine $F_{1,53}$=14.9, p<0.01; and IVSA training session $F_{3,53}$=0.1, p=0.98) in cocaine mice, relative to that in saline mice.

**L**, **M**, Summarized data showing training session-associated differences in the proportions of ALP & $Ca^{2+}$ transient frequency-categorized $Pos_{only}$ neurons (**L**, saline/cocaine × IVSA training session interaction $F_{3,53}$=12.0, p<0.01; saline/cocaine $F_{1,53}$=0.4, p=0.53; and IVSA training session $F_{3,53}$=13.5, p<0.01), and ALP-categorized $Pos_{mixed}$ neurons (**M**, saline/cocaine × IVSA training session interaction $F_{3,53}$=2.0, p=0.12; saline/cocaine $F_{1,53}$=46.3, p<0.01; and IVSA training session $F_{3,53}$=3.1, p=0.04) between saline *vs.* cocaine mice.

**N**, **O**, Summarized data showing training session or the interaction of training session and the IVSA drug-associated differences in the proportions of ALP & $Ca^{2+}$ transient frequency-categorized $Neg_{only}$ neurons (**N**, saline/cocaine × IVSA training session interaction $F_{3,53}$=3.0, p=0.04; saline/cocaine $F_{1,53}$=3.4, p=0.07; and IVSA training session $F_{3,53}$=3.3, p=0.02), and ALP-categorized $Neg_{mixed}$ neurons (**O**, saline/cocaine × IVSA training session interaction $F_{3,53}$=3.1, p=0.03; saline/cocaine $F_{1,53}$=8.6, p<0.01; and IVSA training session $F_{3,53}$=0.9, p=0.45) between saline *vs.* cocaine mice.

Data were analyzed by two-way ANOVA with repeated measures on neuron types and IVSA sessions (**G**, **H**), on IVSA sessions (**I-O**), followed by Bonferroni *post hoc* test. Between different IVSA sessions, *, p<0.05; **, p<0.01. Between saline vs. cocaine, #, p<0.05; ##, 0.01.

### Figure S2

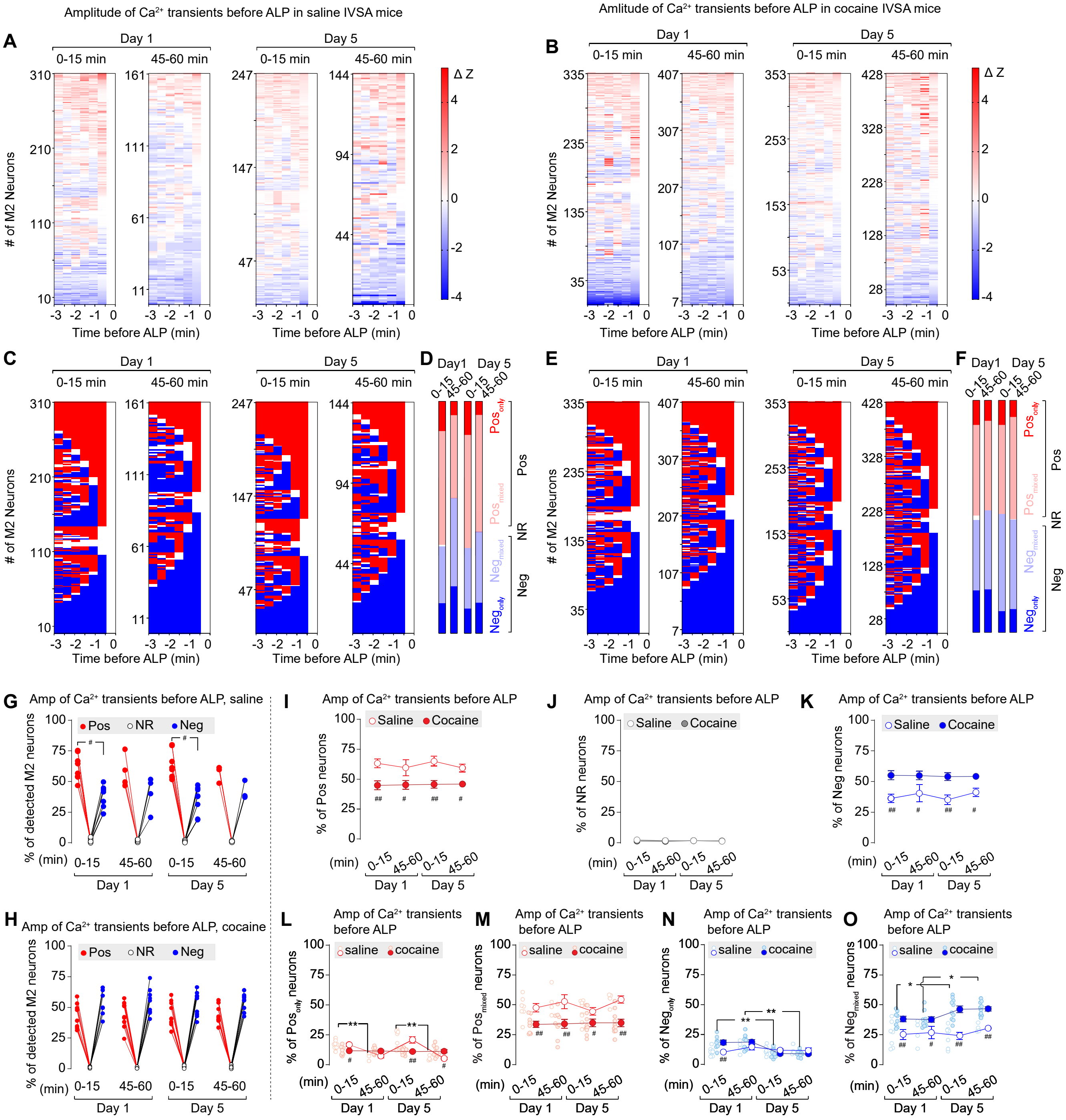

Amplitude of Ca2+ transients before ALP in saline IVSA mice
Amlitude of Ca2+ transients before ALP in cocaine IVSA mice
Day 1
Day 5
0-15 min
45-60 min
### of M2 Neurons
Time before ALP (min)
Δ Z
Pos
NR
Neg
Amp of Ca2+ transients before ALP, saline
Amp of Ca2+ transients before ALP, cocaine
Amp of Ca2+ transients before ALP
% of detected M2 neurons
% of Pos neurons
% of NR neurons
% of Neg neurons
Saline
Cocaine

**Figure S2. Amplitude of $Ca^{2+}$ transients in M2 neurons during 3 min before ALP in saline vs. cocaine mice.**

**A**, **B**, Heat maps of ΔZ scores of $Ca^{2+}$ transient amplitude in detected M2 neurons displayed by rows of six 30-sec blocks within 3 min before ALP in the first (left in each pair of the trace columns) and the last (right in each pair of the trace columns) 15 min during Day 1 and Day 5 IVSA sessions in saline (**A**) and cocaine (**B**) mice.

**C-F**, $Ca^{2+}$ transient amplitude ΔZ based-classification of each 30-sec block of M2 neurons within 3 min before ALP in the first (left in each pair of the trace columns) *vs*. the last (right in each pair of the trace columns) 15 min during Day 1 and Day 5 IVSA sessions in saline (**C**) and cocaine (**D**) mice. The proportions of M2 neurons in each category in a 15-min session are summarized in **E** for saline mice and **F** for cocaine mice.

**G**, **H**, Summarized data showing the percentage of three ALP & $Ca^{2+}$ transient amplitude-based categories of M2 neurons in saline (**G**) and cocaine (**H**) mice. Both saline (**G**, neuron type × IVSA training session interaction $F_{6,40}=0.4$, $p=0.88$; neuron type $F_{1,20}=163.9$, $p<0.01$; and IVSA training session $F_{3,20}=103.8$, $p<0.01$) and cocaine (**H**, neuron type × IVSA training session interaction $F_{6,66}=0.03$, $p>0.99$; neuron type $F_{1,35}=342.2$, $p<0.01$; and IVSA training session $F_{3,33}=384.5$, $p<0.01$) mice demonstrated different proportions among three neuronal types.

**I-K**, Summarized data showing smaller proportions of ALP & $Ca^{2+}$ transient amplitude-categorized Pos neurons (**I**, saline/cocaine × IVSA training session interaction $F_{3,53}=0.3$, $p=0.84$; saline/cocaine $F_{1,53}=34.9$, $p<0.01$; and IVSA training session

$F_{3,53}$=0.3, p=0.86), similar proportions of ALP-categorized NR neurons (**J**, saline/cocaine × IVSA training session interaction $F_{3,53}$=0.8, p=0.50; saline/cocaine $F_{1,53}$=1.2, p=0.27; and IVSA training session $F_{3,53}$=0.7, p=0.58), and larger proportions of ALP-categorized Neg neurons (**K**, saline/cocaine × IVSA training session interaction $F_{3,53}$=0.3, p=0.82; saline/cocaine $F_{1,53}$=36.5, p<0.01; and IVSA training session $F_{3,53}$=0.3, p=0.80) in cocaine mice, relative to that in saline mice.

**L**, **M**, Summarized data showing training session-associated differences in the proportions of ALP & $Ca^{2+}$ transient amplitude-categorized $Pos_{only}$ neurons (**L**, saline/cocaine × IVSA training session interaction $F_{3,53}$=11.4, p<0.01; saline/cocaine $F_{1,53}$=1.1, p=0.30; and IVSA training session $F_{3,53}$=10.9, p<0.01), and ALP-categorized $Pos_{mixed}$ neurons (**M**, saline/cocaine × IVSA training session interaction $F_{3,53}$=0.8, p=0.48; saline/cocaine $F_{1,53}$=35.0, p<0.01; and IVSA training session $F_{3,53}$=0.8, p=0.49) between saline *vs.* cocaine mice.

**N**, **O**, Summarized data showing training session-associated differences in the proportions of ALP & $Ca^{2+}$ transient amplitude-categorized $Neg_{only}$ neurons (**N**, saline/cocaine × IVSA training session interaction $F_{3,53}$=5.7, p<0.01; saline/cocaine $F_{1,53}$=1.8, p=0.18; and IVSA training session $F_{3,53}$=6.5, p<0.01), and ALP-categorized $Neg_{mixed}$ neurons (**O**, saline/cocaine × IVSA training session interaction $F_{3,53}$=1.5, p=0.23; saline/cocaine $F_{1,53}$=52.1, p<0.01; and IVSA training session $F_{3,53}$=2.1, p=0.12) between saline *vs.* cocaine mice.

Data were analyzed by two-way ANOVA with repeated measures on neuron types and IVSA sessions (**G**, **H**), on IVSA sessions (**I-O**), followed by Bonferroni *post hoc*

test. Between different IVSA sessions, *, $p<0.05$; **, $p<0.01$. Between saline vs. cocaine, #, $p<0.05$; ##, 0.01.

#### Figure S3

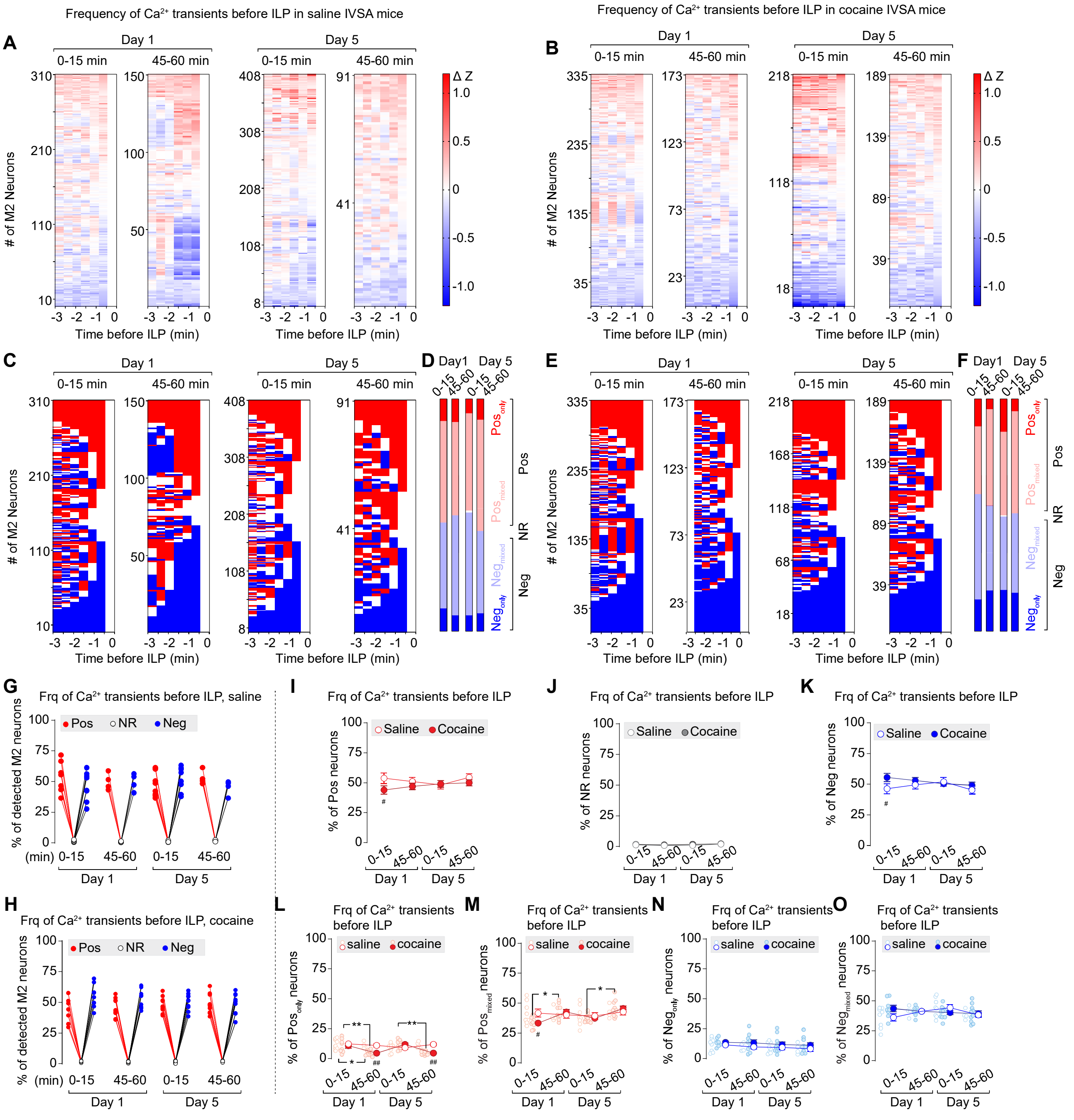

Frequency of Ca2+ transients before ILP in saline IVSA mice
Frequency of Ca2+ transients before ILP in cocaine IVSA mice
A
B
C
D
E
F
G
H
I
J
K
L
M
N
O
Day 1
Day 5
0-15 min
45-60 min
### of M2 Neurons
Time before ILP (min)
Δ Z
Pos
NR
Neg
Frq of Ca2+ transients before ILP, saline
Frq of Ca2+ transients before ILP, cocaine
% of detected M2 neurons
Frq of Ca2+ transients before ILP
Saline
Cocaine
% of Pos neurons
% of NR neurons
% of Neg neurons
% of Pos only neurons
% of Pos mixed neurons
% of Neg only neurons
% of Neg mixed neurons

**S3**. **Frequency of $Ca^{2+}$ transients in M2 neurons during the 3 min before ILP in saline vs. cocaine mice.**

**A**, **B**, Heat maps of ΔZ scores of neuronal $Ca^{2+}$ transient frequency in detected M2 neurons displayed by rows of six 30-sec blocks within 3 min before ILP in the first (left in each pair of the trace columns) and the last (right in each pair of the trace columns) 15 min during Day 1 and Day 5 IVSA sessions in saline (**A**) and cocaine (**B**) mice.

**C-F**, $Ca^{2+}$ transient frequency ΔZ-based classification of **each** 30-sec block of M2 neurons within 3 min before ILP in the first (left in each pair of the trace columns) *vs*. the last (right in each pair of the trace columns) 15 min during Day 1 and Day 5 IVSA sessions in saline (**C**) and cocaine (**E**) mice. The proportions of M2 neurons in each category in a 15-min session is summarized in **D** for saline mice and **F** for cocaine mice.

**G, H**, Summarized data showing the percentage of three ILP & $Ca^{2+}$ transient frequency-based categories of M2 neurons in saline (**G**) and cocaine (**H**) mice. Both saline (**G**, neuron type × IVSA training session interaction $F_{6,40}=0.6$, $p=0.74$; neuron type $F_{1,20}=171.2$, $p<0.01$; and IVSA training session $F_{3,20}=0.0$, $p>0.99$) and cocaine (**H**, neuron type × IVSA training session interaction $F_{6,66}=1.0$, $p=0.40$; neuron type $F_{1,33}=442.0$, $p<0.01$; and IVSA training session $F_{3,33}=150.3$, $p<0.01$) mice demonstrated different proportions among three neuronal types.

**I-K**, Summarized data showing similar proportions of ILP & $Ca^{2+}$ transient frequency-categorized Pos neurons (**I**, saline/cocaine × IVSA training session interaction $F_{3,53}=1.0$, $p=0.40$; saline/cocaine $F_{1,53}=3.2$, $p=0.08$; and IVSA training session

$F_{3,53}$=0.4, p=0.73), ILP-categorized NR neurons (**J**, saline/cocaine × IVSA training session interaction $F_{3,53}$=0.4, p=0.76; saline/cocaine $F_{1,53}$=2.7, p=0.10; and IVSA training session $F_{3,53}$=2.5, p=0.07), and ILP-categorized Neg neurons (**K**, saline/cocaine × IVSA training session interaction $F_{3,53}$=1.1, p=0.36; saline/cocaine $F_{1,53}$=2.5, p=0.12; and IVSA training session $F_{3,53}$=0.6, p=0.60) between saline *vs*. cocaine mice.

**L**, **M**, Summarized data showing significant differences in the proportions of ILP & $Ca^{2+}$ transient frequency-categorized $Pos_{only}$ neurons (**L**, saline/cocaine × IVSA training session interaction $F_{3,53}$=6.0, p<0.01; saline/cocaine $F_{1,53}$=9.8, p<0.01; and IVSA training session $F_{3,53}$=3.4, p=0.03), and ILP-categorized $Pos_{mixed}$ neurons (**M**, saline/cocaine × IVSA training session interaction $F_{3,53}$=2.1, p=0.11; saline/cocaine $F_{1,53}$=0.5, p=0.48; and IVSA training session $F_{3,53}$=2.4, p=0.08) between saline *vs*. cocaine mice.

**N**, **O**, Summarized data showing no detectable differences in the proportions of ILP & $Ca^{2+}$ transient frequency-categorized $Neg_{only}$ neurons (**N**, saline/cocaine × IVSA training session interaction $F_{3,53}$=0.1, p=0.98; saline/cocaine $F_{1,53}$=2.1, p=0.15; and IVSA training session $F_{3,53}$=0.6, p=0.61), or ILP-categorized $Neg_{mixed}$ neurons (**O**, saline/cocaine × IVSA training session interaction $F_{3,53}$=2.1, p=0.11; saline/cocaine $F_{1,53}$=0.4, p=0.55; and IVSA training session $F_{3,53}$=0.7, p=0.53) between saline *vs*. cocaine mice.

Data were analyzed by two-way ANOVA with repeated measures on neuron types and IVSA sessions (**G**, **H**), on IVSA sessions (**I-O**), followed by Bonferroni *post hoc*

test. Between different IVSA sessions, *, $p<0.05$; **, $p<0.01$. Between saline vs. cocaine, #, $p<0.05$; ##, 0.01.

#### Figure S4

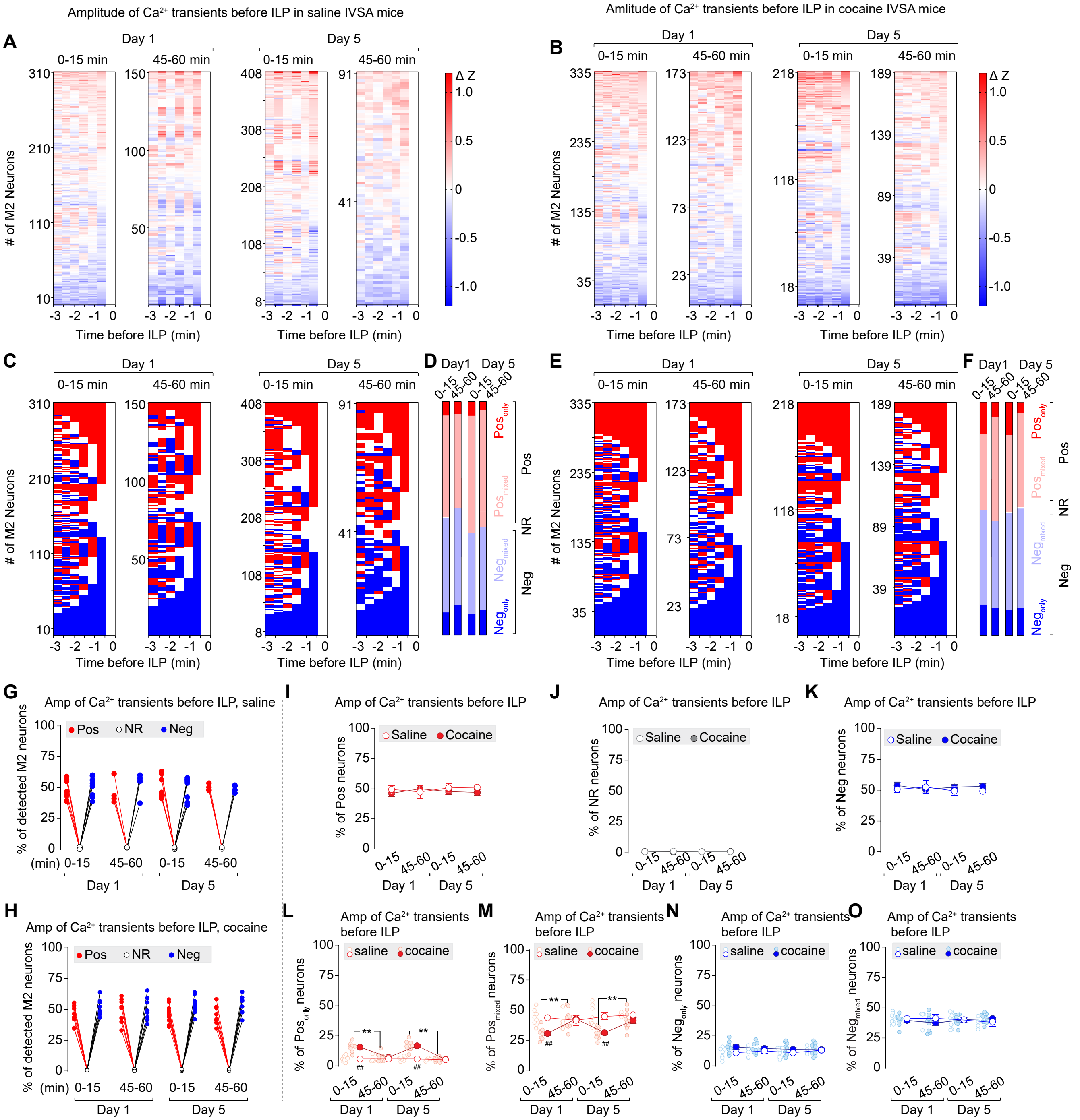

Amplitude of Ca2+ transients before ILP in saline IVSA mice
Amlitude of Ca2+ transients before ILP in cocaine IVSA mice
A
B
C
D
E
F
G
H
I
J
K
L
M
N
O
Day 1
Day 5
0-15 min
45-60 min
### of M2 Neurons
Time before ILP (min)
Δ Z
Pos
NR
Neg
Amp of Ca2+ transients before ILP, saline
Amp of Ca2+ transients before ILP, cocaine
Amp of Ca2+ transients before ILP
% of detected M2 neurons
% of Pos neurons
% of NR neurons
% of Neg neurons
Saline
Cocaine
(min)
Amp of Ca2+ transients before ILP
% of Pos only neurons
% of Pos mixed neurons
% of Neg only neurons
% of Neg mixed neurons
saline
cocaine

**Figure S4**. **Amplitude of $Ca^{2+}$ transients in M2 neurons during the 3 min before ILP in saline vs. cocaine mice.**

**A**, **B**, Heat maps of ΔZ scores of neuronal $Ca^{2+}$ transient amplitude in detected M2 neurons displayed by rows of six 30-sec blocks within 3 min before ILP in the first (left in each pair of the trace columns) and the last (right in each pair of the trace columns) 15 min during Day 1 and Day 5 IVSA sessions in saline (**A**) and cocaine (**B**) mice.

**C-F**, $Ca^{2+}$ transient amplitude ΔZ-based classification of **each** 30-sec block of M2 neurons within 3 min before ILP in the first (left in each pair of the trace columns) *vs*. the last (right in each pair of the trace columns) 15 min during Day 1 and Day 5 IVSA sessions in saline (**C**) and cocaine (**E**) mice. The proportions of M2 neurons in each category in a 15-min session is summarized in **D** for saline mice and **F** for cocaine mice.

**G, H**, Summarized data showing the percentage of three ILP & $Ca^{2+}$ transient amplitude-based categories of M2 neurons in saline (**G**) and cocaine (**H**) mice. Both saline (**G**, neuron type × IVSA training session interaction $F_{6,40}=0.2$, p=0.98; neuron type $F_{1,20}=240.5$, p<0.01; and IVSA training session $F_{3,20}=1.9$, p=0.17) and cocaine (**H**, neuron type × IVSA training session interaction $F_{6,66}=0.3$, p=0.93; neuron type $F_{1,33}=480.3$, p<0.01; and IVSA training session $F_{3,33}=0.4$, p=0.74) mice demonstrated different proportions among three neuronal types.

**I-K**, Summarized data showing similar proportions of ILP & $Ca^{2+}$ transient amplitude-categorized Pos neurons (**I**, saline/cocaine × IVSA training session interaction $F_{3,53}=0.4$, p=0.76; saline/cocaine $F_{1,53}=0.8$, p=0.38; and IVSA training session

$F_{3,53}$=0.1, p=0.96), ILP-categorized NR neurons (**J**, saline/cocaine × IVSA training session interaction $F_{3,53}$=0.8, p=0.49; saline/cocaine $F_{1,53}$=1.2, p=0.28; and IVSA training session $F_{3,53}$=0.1, p=0.98), and ILP-categorized Neg neurons (**K**, saline/cocaine × IVSA training session interaction $F_{3,53}$=0.3, p=0.80; saline/cocaine $F_{1,53}$=1.0, p=0.32; and IVSA training session $F_{3,53}$=0.1, p=0.96) between saline *vs*. cocaine mice.

**L**, **M**, Summarized data showing significant differences in the proportions of ILP & $Ca^{2+}$ transient amplitude-categorized $Pos_{only}$ neurons (**L**, saline/cocaine × IVSA training session interaction $F_{3,53}$=12.0, p<0.01; saline/cocaine $F_{1,53}$=49.2, p<0.01; and IVSA training session $F_{3,53}$=14.2, p<0.01), and ILP-categorized $Pos_{mixed}$ neurons (**M**, saline/cocaine × IVSA training session interaction $F_{3,53}$=3.6, p=0.02; saline/cocaine $F_{1,53}$=17.0, p<0.01; and IVSA training session $F_{3,53}$=2.9, p=0.04) between saline *vs*. cocaine mice.

**N**, **O**, Summarized data showing no detectable differences in the proportions of ILP & $Ca^{2+}$ transient amplitude-categorized $Neg_{only}$ neurons (**N**, saline/cocaine × IVSA training session interaction $F_{3,53}$=0.6, p=0.65; saline/cocaine $F_{1,53}$=3.3, p=0.07; and IVSA training session $F_{3,53}$=0.2, p=0.88), or ILP-categorized $Neg_{mixed}$ neurons (**O**, saline/cocaine × IVSA training session interaction $F_{3,53}$=1.2, p=0.33; saline/cocaine $F_{1,53}$=0.1, p=0.80; and IVSA training session $F_{3,53}$=0.1, p=0.94) between saline *vs*. cocaine mice.

Data were analyzed by two-way ANOVA with repeated measures on neuron types and IVSA sessions (**G**, **H**), on IVSA sessions (**I-O**), followed by Bonferroni *post hoc*

test. Between different IVSA sessions, *, $p<0.05$; **, $p<0.01$. Between saline vs. cocaine, #, $p<0.05$; ##, 0.01.

#### Figure S5

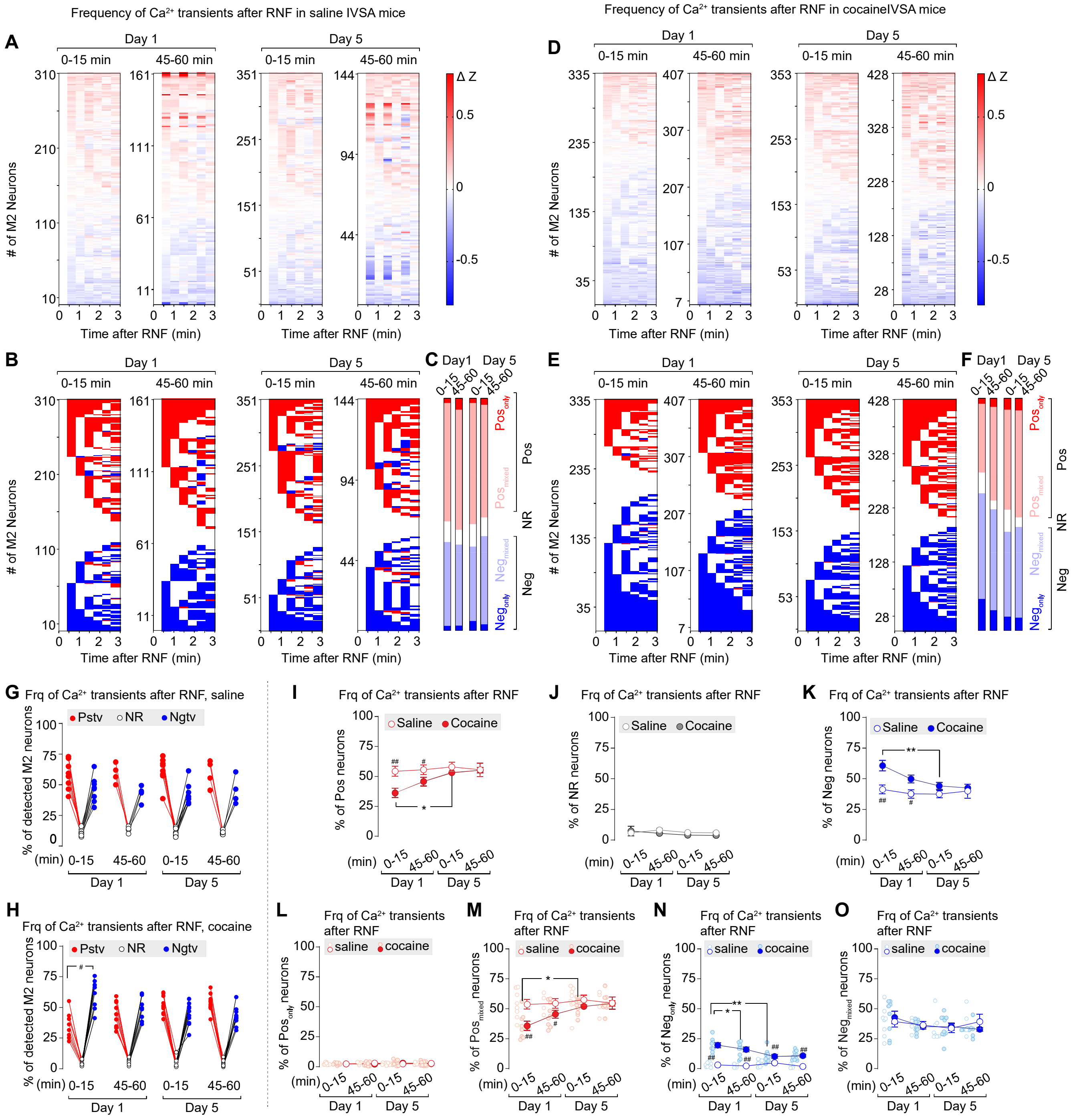

Frequency of Ca2+ transients after RNF in saline IVSA mice
Frequency of Ca2+ transients after RNF in cocaineIVSA mice
A
B
C
D
E
F
Day 1
Day 5
0-15 min
45-60 min
### of M2 Neurons
Time after RNF (min)
Δ Z
Pos
NR
Neg
G
Frq of Ca2+ transients after RNF, saline
Pstv
NR
Ngtv
% of detected M2 neurons
H
Frq of Ca2+ transients after RNF, cocaine
I
Frq of Ca2+ transients after RNF
Saline
Cocaine
% of Pos neurons
J
% of NR neurons
K
% of Neg neurons
L
Frq of Ca2+ transients after RNF
saline
cocaine
M
N
O
(min)

**Figure S5**. **Frequency of $Ca^{2+}$ transients in M2 neurons during the 3 min after RNF in saline vs. cocaine mice.**

**A**, **B**, Heat maps of ΔZ scores of neuronal $Ca^{2+}$ transient frequency in detected M2 neurons displayed by rows of six 30-sec blocks within 3 min after RNF in the first (left in each pair of the trace columns) and the last (right in each pair of the trace columns) 15 min during Day 1 and Day 5 IVSA sessions in saline (**A**) and cocaine (**B**) mice.

**C-F**, $Ca^{2+}$ transient frequency ΔZ-based classification of **each** 30-sec block of M2 neuron within 3 min after RNF in the first (left in each pair of the trace columns) *vs*. the last (right in each pair of the trace columns) 15 min during Day 1 and Day 5 IVSA sessions in saline (**C**) and cocaine (**E**) mice. The proportions of M2 neurons in each category in a 15-min session are summarized in **D** for saline mice and **F** for cocaine mice.

**G**, **H**, Summarized data showing the percentage of three RNF & $Ca^{2+}$ transient frequency-based categories of M2 neurons in saline (**G**) and cocaine (**H**) mice. Both saline (**G**, neuron type × IVSA training session interaction $F_{6,40}=0.2$, $p=0.97$; neuron type $F_{1,23}=117$, $p<0.01$; and IVSA training session $F_{3,20}=231$, $p<0.01$) and cocaine (**H**, neuron type × IVSA training session interaction $F_{6,66}=6.7$, $p<0.01$; neuron type $F_{1,39}=258$, $p<0.01$; and IVSA training session $F_{3,33}=85.0$, $p<0.01$) mice demonstrated different proportions among three neuronal types.

**I-K**, Summarized data showing lower proportions of RNF & $Ca^{2+}$ transient frequency-categorized Pos neurons (**I**, saline/cocaine × IVSA training session interaction $F_{3,53}=2.1$, $p=0.11$; saline/cocaine $F_{1,53}=9.3$, $p=0.04$; and IVSA training session

$F_{3,53}$=3.7, p=0.02), similar proportions of RNF & $Ca^{2+}$ transient frequency-categorized NR neurons (**J**, saline/cocaine × IVSA training session interaction $F_{3,53}$=0.6, p=0.65; saline/cocaine $F_{1,53}$=1.0, p=0.31; and IVSA training session $F_{3,53}$=0.7, p=0.58), and higher proportions of RNF & $Ca^{2+}$ transient frequency-categorized Neg neurons (**K**, saline/cocaine × IVSA training session interaction $F_{3,53}$=2.1, p=0.11; saline/cocaine $F_{1,53}$=15.7, p<0.01; and IVSA training session $F_{3,53}$=3.8, p=0.02) in cocaine mice, relative to that in saline mice.

**L**, **M**, Summarized data showing training session-associated differences in the proportions of RNF & $Ca^{2+}$ transient frequency -categorized $Pos_{only}$ neurons (**L**, saline/cocaine × IVSA training session interaction $F_{3,53}$=0.5, p=0.68; saline/cocaine $F_{1,53}$=0.1, p=0.80; and IVSA training session $F_{3,53}$=0.2, p=0.93), and RNF-categorized $Pos_{mixed}$ neurons (**M**, saline/cocaine × IVSA training session interaction $F_{3,53}$=2.2, p=0.10; saline/cocaine $F_{1,53}$=10.0, p<0.01; and IVSA training session $F_{3,53}$=3.8, p=0.02) between saline *vs.* cocaine mice.

**N**, **O**, Summarized data showing training session-associated differences in the proportions of RNF & $Ca^{2+}$ transient frequency-categorized $Neg_{only}$ neurons (**N**, saline/cocaine × IVSA training session interaction $F_{3,53}$=5.2, p<0.01; saline/cocaine $F_{1,53}$=91.0, p<0.01; and IVSA training session $F_{3,53}$=3.7, p=0.02), and RNF-categorized $Neg_{mixed}$ neurons (**O**, saline/cocaine × IVSA training session interaction $F_{3,53}$=0.6, p=0.61; saline/cocaine $F_{1,53}$=0.1, p=0.80; and IVSA training session $F_{3,53}$=1.4, p=0.27) between saline *vs.* cocaine mice.

Data were analyzed by two-way ANOVA with repeated measures on neuron types and IVSA sessions (**G**, **H**), on IVSA sessions (**I-O**), followed by Bonferroni's *post hoc*

test. Between different IVSA sessions, *, $p<0.05$; **, $p<0.01$. Between saline vs. cocaine, #, $p<0.05$; ##, 0.01.

#### Figure S6

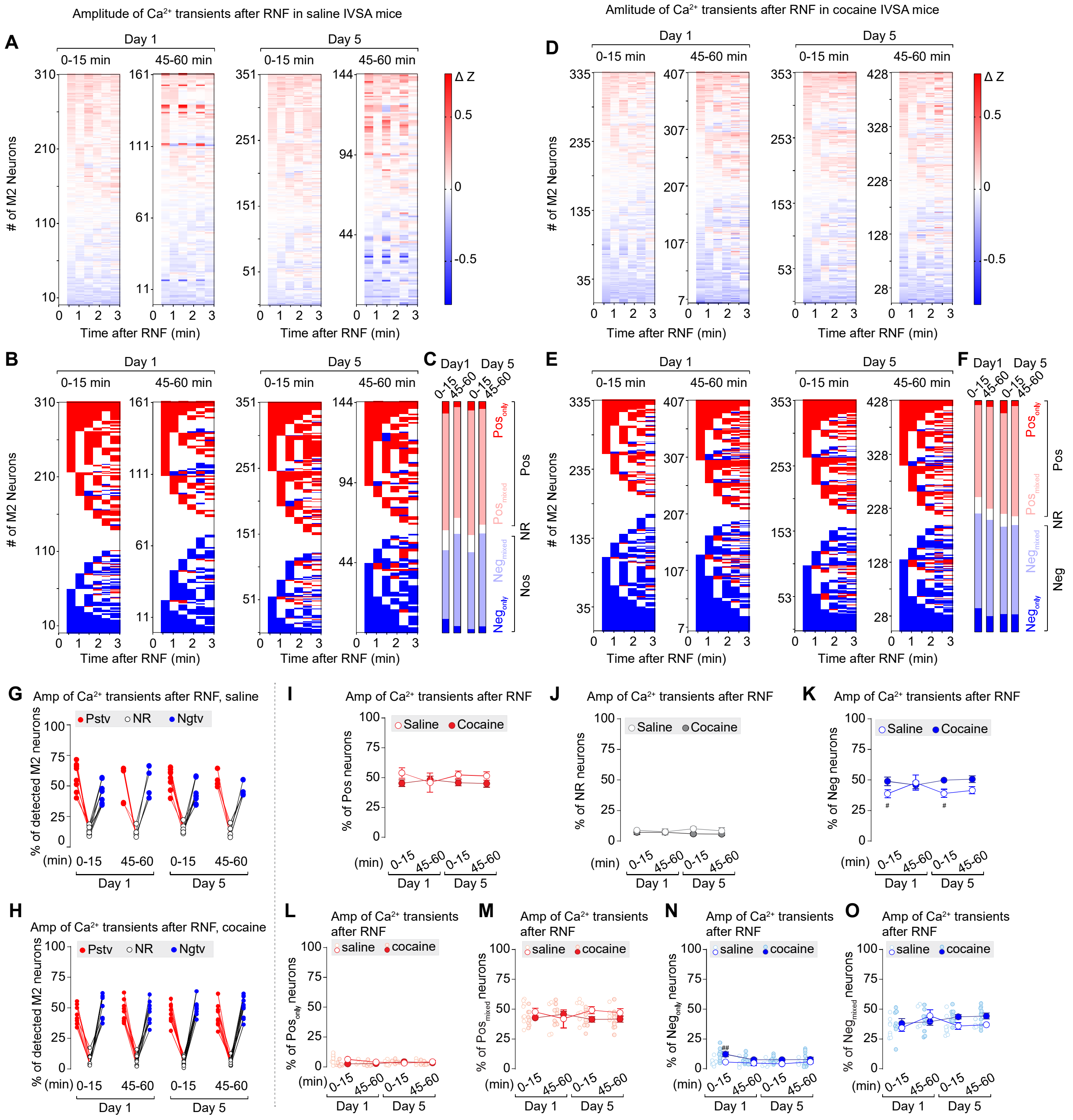

Amplitude of Ca2+ transients after RNF in saline IVSA mice
Amlitude of Ca2+ transients after RNF in cocaine IVSA mice
A
B
C
D
E
F
G
H
I
J
K
L
M
N
O
Day 1
Day 5
0-15 min
45-60 min
### of M2 Neurons
Time after RNF (min)
Δ Z
Pos only
Pos mixed
Neg mixed
Neg only
Pos
NR
Neg
Amp of Ca2+ transients after RNF, saline
Amp of Ca2+ transients after RNF, cocaine
Pstv
NR
Ngtv
% of detected M2 neurons
Amp of Ca2+ transients after RNF
Saline
Cocaine
% of Pos neurons
% of NR neurons
% of Neg neurons
% of Pos only neurons
% of Pos mixed neurons
% of Neg only neurons
% of Neg mixed neurons
(min)

**Figure S6**. **Amplitude of $Ca^{2+}$ transients in M2 neurons during the 3 min after RNF in saline vs. cocaine mice.**

**A**, **B**, Heat maps of ΔZ scores of neuronal $Ca^{2+}$ transient amplitude in detected M2 neurons displayed by rows of six 30-sec blocks within 3 min after RNF in the first (left in each pair of the trace columns) and the last (right in each pair of the trace columns) 15 min during Day 1 and Day 5 IVSA sessions in saline (**A**) and cocaine (**B**) mice.

**C-F**, $Ca^{2+}$ transient amplitude ΔZ-based classification of **each** 30-sec block of M2 neuron within 3 min after RNF in the first (left in each pair of the trace columns) *vs*. the last (right in each pair of the trace columns) 15 min during Day 1 and Day 5 IVSA sessions in saline (**C**) and cocaine (**E**) mice. The proportions of M2 neurons in each category in a 15-min session are summarized in **D** for saline mice and **F** for cocaine mice.

**G**, **H**, Summarized data showing the percentage of three RNF & $Ca^{2+}$ transient amplitude-based categories of M2 neurons in saline (**G**) and cocaine (**H**) mice. Both saline (**G**, neuron type × IVSA training session interaction $F_{6,40}=0.7$, $p=0.67$; neuron type $F_{1,23}=90.0$, $p<0.01$; and IVSA training session $F_{3,20}=42.2$, $p<0.01$) and cocaine (**H**, neuron type × IVSA training session interaction $F_{6,66}=0.4$, $p=0.87$; neuron type $F_{1,39}=254.3$, $p<0.01$; and IVSA training session $F_{3,33}=140.8$, $p<0.01$) mice demonstrated different proportions among three neuronal types.

**I-K**, Summarized data showing the proportions of RNF & $Ca^{2+}$ transient amplitude-categorized Pos neurons (**I**, saline/cocaine × IVSA training session interaction $F_{3,53}=0.8$, $p=0.51$; saline/cocaine $F_{1,53}=3.4$, $p=0.07$; and IVSA training session

$F_{3,53}$=0.2, p=0.92), RNF & $Ca^{2+}$ transient amplitude-categorized NR neurons (**J**, saline/cocaine × IVSA training session interaction $F_{3,53}$=0.6, p=0.62; saline/cocaine $F_{1,53}$=4.3, p=0.04; and IVSA training session $F_{3,53}$=0.2, p=0.92), and RNF & $Ca^{2+}$ transient amplitude-categorized Neg neurons (**K**, saline/cocaine × IVSA training session interaction $F_{3,53}$=1.4, p=0.25; saline/cocaine $F_{1,53}$=8.5, p=0.01; and IVSA training session $F_{3,53}$=0.3, p=0.79) in cocaine mice, relative to that in saline mice.

**L**, **M**, Summarized data showing training session-associated differences in the proportions of RNF & $Ca^{2+}$ transient amplitude -categorized $Pos_{only}$ neurons (**L**, saline/cocaine × IVSA training session interaction $F_{3,53}$=4.0, p=0.01; saline/cocaine $F_{1,53}$=4.6, p=0.04; and IVSA training session $F_{3,53}$=0.7, p=0.58), and RNF-categorized $Pos_{mixed}$ neurons (**M**, saline/cocaine × IVSA training session interaction $F_{3,53}$=1.1, p=0.36; saline/cocaine $F_{1,53}$=2.5, p=0.12; and IVSA training session $F_{3,53}$=0.1, p=0.96) between saline *vs.* cocaine mice.

**N**, **O**, Summarized data showing training session-associated differences in the proportions of RNF & $Ca^{2+}$ transient amplitude-categorized $Neg_{only}$ neurons (**N**, saline/cocaine × IVSA training session interaction $F_{3,53}$=0.9, p=0.46; saline/cocaine $F_{1,53}$=11.3, p<0.01; and IVSA training session $F_{3,53}$=1.7, p=0.18), and RNF-categorized $Neg_{mixed}$ neurons (**O**, saline/cocaine × IVSA training session interaction $F_{3,53}$=0.2, p=0.89; saline/cocaine $F_{1,53}$=4.4, p=0.04; and IVSA training session $F_{3,53}$=1.8, p=0.161) between saline *vs.* cocaine mice.

Data were analyzed by two-way ANOVA with repeated measures on neuron types and IVSA sessions (**G**, **H**), on IVSA sessions (**I-O**), followed by Bonferroni's *post hoc*

test. Between different IVSA sessions, *, $p<0.05$; **, $p<0.01$. Between saline vs. cocaine, #, $p<0.05$; ##, 0.01.

#### Figure S7

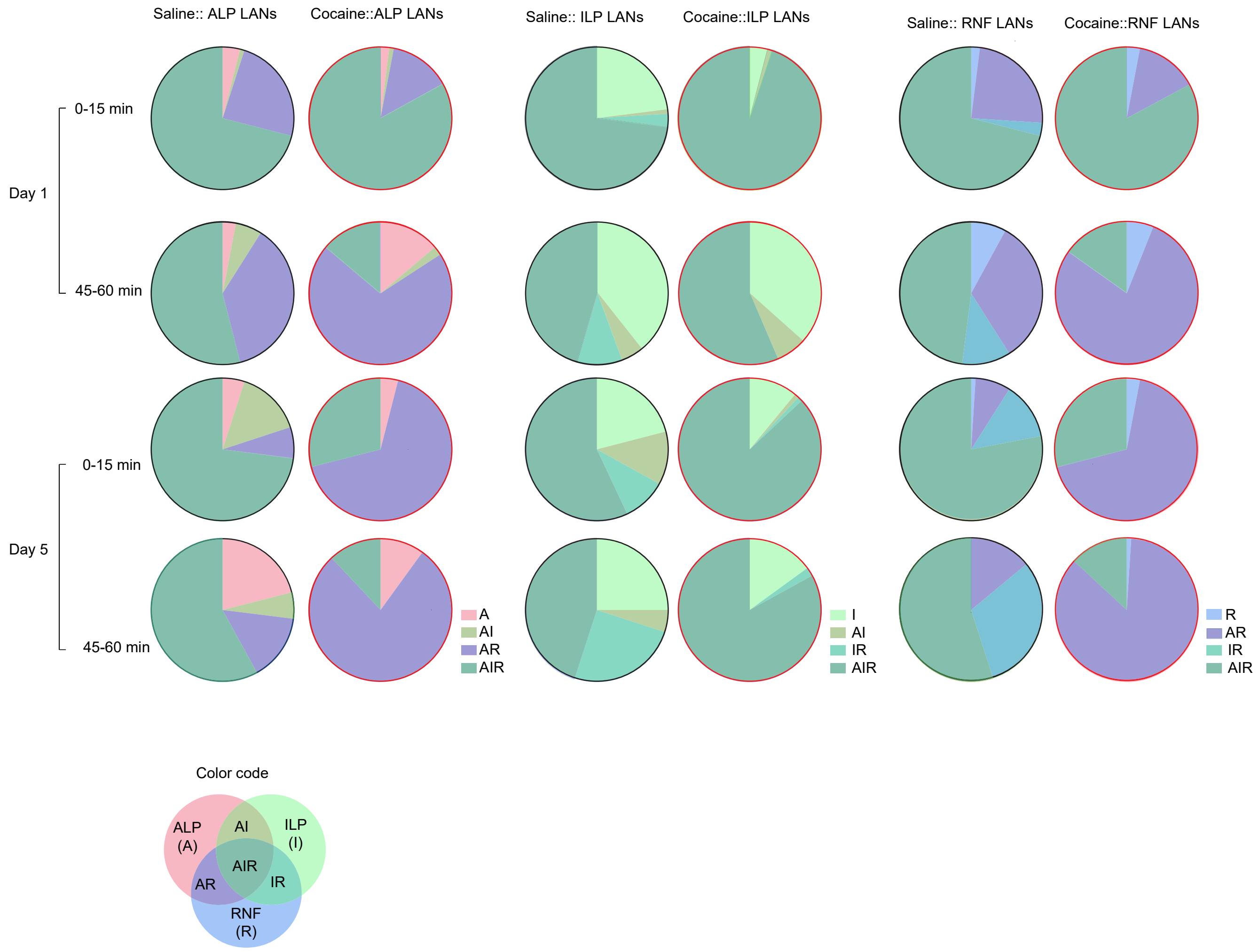
Saline:: ALP LANs
Cocaine::ALP LANs
Saline:: ILP LANs
Cocaine::ILP LANs
Saline:: RNF LANs
Cocaine::RNF LANs
Day 1
0-15 min
45-60 min
Day 5
0-15 min
45-60 min
A
AI
AR
AIR
I
AI
IR
AIR
R
AR
IR
AIR
Color code
ALP
(A)
AI
ILP
(I)
AIR
AR
IR
RNF
(R)

**Figure S7**. Pie charts showing the **overlaps of IVSA event-associated LANs in M2.**

**Table S1. 3-way ANOVA analysis of neuronal $Ca^{2+}$ transient in M2**

| | | Total $Ca^{2+}$ transients | Frequency | Amplitude |
|---|---|---|---|---|
| | Figures | **Figs. 2C, D, I** | **Figs. 2E, F, J** | **Figs. 2G, H, K** |
| Factor 1 | 0-15 vs. 45-60 min ("min") | $F(1, 3337) = 2217$, $p<0.0001$ | $F(1, 3337) = 1675$, $p<0.0001$ | $F(1, 3337) = 400.6$, $p<0.0001$ |
| Factor 2 | saline vs. cocaine ("S/C") | $F(1, 3337) = 887.2$, $p<0.0001$ | $F(1, 3337) = 466.2$, $p<0.0001$ | $F(1, 3337) = 306.5$, $p<0.0001$ |
| Factor 3 | Day 1 vs. 5 (D) | $F(1, 3337) = 0.5720$, $p=0.4495$ | $F(1, 3337) = 103.0$, $p<0.0001$ | $F(1, 3337) = 69.07$, $p<0.0001$ |
| Interactions | min x S/C | $F(1, 3337) = 427.4$, $p<0.0001$ | $F(1, 3337) = 210.1$, $p<0.0001$ | $F(1, 3337) = 116.3$, $p<0.0001$ |
| | min x D | $F(1, 3337) = 1201$, $p<0.0001$ | $F(1, 3337) = 1541$, $p<0.0001$ | $F(1, 3337) = 105.3$, $p<0.0001$ |
| | S/C x D | $F(1, 3337) = 252.7$, $p<0.0001$ | $F(1, 3337) = 205.4$, $p<0.0001$ | $F(1, 3337) = 73.32$, $p<0.0001$ |
| | min x S/C x D | $F(1, 3337) = 75.75$, $p<0.0001$ | $F(1, 3337) = 92.88$, $p<0.0001$ | $F(1, 3337) = 14.02$, $p=0.0002$ |

**Bonferroni's post hoc test**

| | | Total $Ca^{2+}$ transients | Frequency | Amplitude |
|---|---|---|---|---|
| | Figures | **Figs. 2C, D, I** | **Figs. 2E, F, J** | **Figs. 2G, H, K** |
| within saline mice | 0-15 min on Day 1 vs. Day 5 | <0.0001 | <0.0001 | <0.0001 |
| | 45-60 min on Day 1 vs. Day 5 | >0.9999 | >0.9999 | >0.9999 |
| | 0-15 vs. 45-60 min on Day 1 | <0.0001 | <0.0001 | >0.9999 |
| | 0-15 vs. 45-60 min on Day 5 | <0.0001 | <0.0001 | 0.008 |
| within cocaine mice | 0-15 min on Day 1 vs. Day 5 | <0.0001 | <0.0001 | <0.0001 |
| | 45-60 min on Day 1 vs. Day 5 | <0.0001 | <0.0001 | <0.0001 |
| | 0-15 vs. 45-60 min on Day 1 | <0.0001 | <0.0001 | <0.0001 |
| | 0-15 vs. 45-60 min on Day 5 | <0.0001 | 0.3456 | 0.0641 |
| saline vs. cocaine mice | 0-15 min on Day 1 | <0.0001 | <0.0001 | 0.7279 |
| | 45-60 min on Day 1 | <0.0001 | <0.0001 | <0.0001 |
| | 0-15 min on Day 5 | <0.0001 | <0.0001 | <0.0001 |
| | 45-60 min on Day 5 | <0.0001 | <0.0001 | <0.0001 |
